# Atf3 Integrates Lipid and Cytoskeletal Remodeling to Drive Macrophage Fusion

**DOI:** 10.64898/2026.04.01.715652

**Authors:** Andreia Correia, Naziha Mroueh, Esther Katharina Wollert, Dimitrije Stankovic, Gábor Csordás, Christian Jüngst, Antonio García-Bernardo Tartiere, Martim Dias Gomes, Sandra Iden, Mirka Uhlirova

## Abstract

Macrophages are highly plastic innate immune cells that can form multinucleated giant cells (MGCs) under physiological and pathological conditions, such as osteoclasts and foreign body giant cells. The mechanisms governing macrophage multinucleation remain incompletely understood. Here, we identify activating transcription factor 3 (Atf3) as an essential regulator of MGC formation via cell-cell fusion in response to persistent Interleukin 4 (IL-4) and STAT6 signaling, characteristic of the foreign body reaction. *Atf3*-deficient macrophages activate STAT6-dependent transcriptional programs in response to IL-4, including fusion-associated genes, yet fail to undergo multinucleation. This defect is associated with impaired actin cytoskeleton remodeling, abnormal nuclear morphology, reduced lamin A/C expression, and genome instability. Mechanistically, loss of *Atf3* derepresses the *Cholesterol-25-hydroxylase* (Ch25h), elevating 25-hydroxycholesterol (25-HC), suppressing the mevalonate pathway, and reducing cholesterol and isoprenoid biosynthesis essential for cytoskeletal dynamics. Deletion of *Ch25h* in *Atf3*-deficient macrophages restores cholesterol levels and actin turnover, but not lamin A/C or fusion. These findings establish Atf3 as a central transcriptional node integrating lipid metabolism with cytoskeletal and nucleoskeletal remodeling to control IL-4-driven macrophage multinucleation.

## INTRODUCTION

Macrophages are versatile sentinels of the innate immune system, serving as professional phagocytes, antigen-presenting cells and coordinators of tissue repair (Mass *et al*, 2023; Park *et al*, 2022; Wynn & Vannella, 2016). A remarkable feature of macrophages is their ability to form multinucleated giant cells (MGCs) under specific physiological and pathological conditions. Osteoclasts represent the prototypical physiological MGCs that differentiate in response to the receptor activator of NF-κB ligand (RANKL) to mediate bone remodeling and homeostasis (Levaot *et al*, 2015; Yasuda *et al*, 1998). In contrast, pathological MGCs arise during chronic inflammation associated with granuloma formation in infectious, autoinflammatory, and allergic contexts. Prominent examples include Langhans cells in tuberculosis, MGCs found in giant cell arteritis (GCA) and atherosclerosis, and foreign body giant cells (FBGCs) formed in response to implanted biomaterials or tissue-damaging amyloid fibrils (Ahmadzadeh *et al*, 2022; Anderson *et al*, 2008; Bodin *et al*, 2010; Brooks *et al*, 2019; Samokhin *et al*, 2010). Importantly, the differentiation of distinct MGC subtypes is dictated by the local inflammatory milieu. For example, Langhans cells form in response to Toll-like receptor 2 (TLR2) and tumor necrosis factor (TNF) whereas FBGCs are driven by a Th2-polarized environment rich in IL-4 and IL-13, highlighting the existence of context-dependent transcriptional programs governing MGC differentiation (Helming & Gordon, 2009; Herrtwich *et al*, 2016; McInnes & Rennick, 1988; McNally & Anderson, 1995; Puissegur *et al*, 2007).

Macrophage multinucleation occurs via two primary mechanisms: cell-cell fusion of individual macrophages, and acytokinetic cell division, where nuclear division proceeds without cytokinesis (Helming & Gordon, 2009; Herrtwich *et al*., 2016; Takegahara *et al*, 2016). Both processes require tight coordination of transcriptional reprograming, cytoskeletal and membrane remodeling. The process of cell fusion is particularly complex, involving migration, acquisition of a fusion-competent state, assembly of adhesive, force-generating podosomes, and finally the formation of the fusion pore and plasma membrane merging (Dufrancais *et al*, 2021; Helming & Gordon, 2009). These steps critically depend on actin cytoskeleton dynamics, as disrupting key regulators like WASP, Rac-1, Cdc42, Filamin A or the Arp2/3 complex, severely impairs fusion efficiency (Arya *et al*, 2021; DeFife *et al*, 1997; Faust *et al*, 2019; Gonzalo *et al*, 2010; Jay *et al*, 2007; Leung *et al*, 2010). Concurrently, membrane lipid composition, particularly cholesterol content, provides another layer of regulation by modulating membrane fluidity and protein distribution, thereby affecting the organization of the cytoskeleton (Brukman *et al*, 2019; Lee & Bensinger, 2022; Lösslein *et al*, 2021). However, how macrophages integrate extracellular signals with these intrinsic programs of lipid metabolism, cytoskeletal rearrangement, and transcriptional control to drive multinucleation remains incompletely understood.

The stress-responsive Activating transcription factor 3 (Atf3), a member of the ATF/CREB family, has emerged as a key regulator of inflammatory and metabolic programs (Hu *et al*, 2024; Jadhav & Zhang, 2017; Rynes *et al*, 2012). In macrophages, Atf3 attenuates pro-inflammatory cytokine expression driven by Toll-like receptor (TLR) signaling (De Nardo *et al*, 2014; Gilchrist *et al*, 2006; Labzin *et al*, 2015) while also suppressing the transcription of *Cholesterol-25-hydroxylase* (*Ch25h*) (Gold *et al*, 2012). Ch25h is an ER-resident enzyme that produces 25-hydroxycholesterol (25-HC) (Lund *et al*, 1998), a potent oxysterol that restricts cholesterol and isoprenoid synthesis via the mevalonate pathway and alters plasma membrane cholesterol content (Griffiths & Wang, 2022). Furthermore, Atf3 promotes macrophage migration and metabolic phenotypes associated with alternative activation, which are processes permissive for MGC formation (Hu *et al*., 2024; Lösslein *et al*., 2021). The physiological importance of Atf3 is highlighted by the increased susceptibility of Atf3-deficient mice to septic shock and atherosclerosis (De Nardo *et al*., 2014; Gilchrist *et al*., 2006; Gold *et al*., 2012). Intriguingly, emerging evidence also links Atf3 to regulation of cytoarchitecture, including components of the cytoskeleton and nucleoskeleton (Boespflug *et al*, 2014; Donohoe *et al*, 2018; Du *et al*, 2022; Sekyrova *et al*, 2010). Together, these functions position Atf3 as a potential integrator of the inflammatory, metabolic, and structural remodeling required for macrophage multinucleation.

Here, we identify Atf3 as a central regulator of IL-4-driven macrophage fusion. We demonstrate that *Atf3* deficiency leads to the derepression of *Ch25h*, resulting in 25-HC accumulation and suppression of the mevalonate pathway. These metabolic changes cause profound defects in membrane and actin dynamics, leading to aberrant morphology and impaired multinucleation. Remarkably, while restoring cholesterol and isoprenoid synthesis rescues cellular morphology, it fails to rescue fusion, uncovering a distinct, Ch25h-independent role for Atf3 in maintaining nuclear lamina integrity. Together, our findings establish Atf3 as a critical transcriptional hub that couples inflammatory signaling with metabolic, cytoskeletal, and nuclear programs essential for macrophage multinucleation.

## RESULTS

### Atf3 is required for macrophage multinucleation downstream of IL-4

Macrophage multinucleation occurs physiologically in the bone, where osteoclasts form in response to RANKL, and pathologically during chronic inflammation such as *Mycobacteria* infection or foreign body reactions driven by TLR2 and IL-4/IL-13 signaling, respectively. To determine whether Atf3 is involved in this process, we assessed its expression in mouse bone marrow-derived macrophages (BMDMs) exposed to different MGC-inducing stimuli, including RANKL, TLR2 agonists and IL-4. In this study, multinucleation is defined as the presence of more than two nuclei within a single macrophage, a criterion used throughout to define multinucleated giant cells (MGCs). Atf3 mRNA and protein levels were significantly induced after one day of IL-4 treatment but remained unchanged in response to RANKL or the synthetic TLR2 agonists, FSL-1 and Pam3CSK4 (Fig. 1A, B). In *control* BMDM cultures, giant cells appeared after one day of IL-4 treatment and increased in size and number over time (Fig. 1P-Q). Strikingly, macrophages isolated from whole-body *Atf3* knockout (*Atf3*^*KO*^) mice generated by CRISPR/Cas9 (Fig. 1B, C) failed to multinucleate in response to IL-4 (Fig. 1D-O, Q). Increasing cell density, IL-4 concentration, or both, or culturing cells on a fusogenic Permanox substrate (Helming & Gordon, 2009), did not mitigate the multinucleation defect of *Atf3*^*KO*^ cells (Fig. 1r-u). The absence of MGCs was also evident in IL-4-stimulated macrophages isolated from the peritoneal cavity of *Atf3*-deficient mice (Fig. 1V-X), indicating that Atf3 is essential for IL-4-driven macrophage multinucleation.

**Figure 1.**
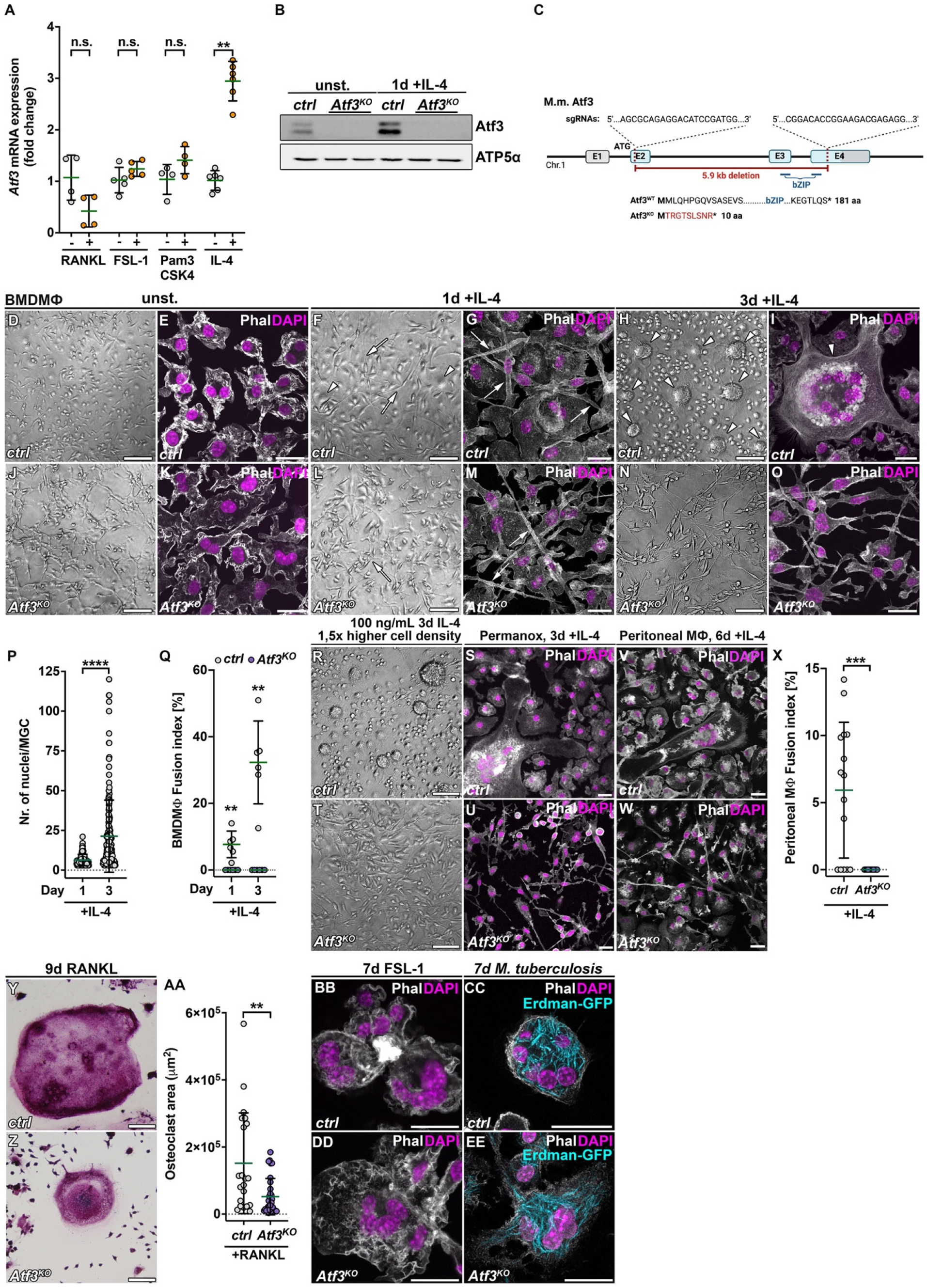
Atf3 is required for macrophage multinucleation downstream of IL-4. **(A)** *Atf3* mRNA expression in *control (ctrl)* BMDMs upon one day of RANKL, FSL-1, Pam3CSK4 and IL-4 stimulation (+), relative to unstimulated (-) *control* cells. **(B)** Western blot analysis revealed induction of Atf3 protein upon IL-4 stimulation. Atf3 protein is undetectable in BMDMs isolated from *Atf3*^*KO*^ mice relative to *control* macrophages. ATP5α was used as a loading control. **(C)** Schematic representation of *Atf3* locus targeted by two sgRNAs to generate a deletion of 5933 bp, including the bZIP domain. Boxes represent the four exons (E1-E4), which are interspaced by introns. The start codon (ATG) is in the second exon. Grey areas of E1 and E4 represent 5’- and 3’-UTR sequences. **(D-O)** Representative bright field (D, F, H, J, L, N) and confocal (E, G, I, K, M, O) micrographs of *control* (D-I) and *Atf3*^*KO*^ (J-O) BMDMs unstimulated (D, E, J, K) and stimulated with IL-4 for one day (F, G, L, M) and three days (H, I, N, O). Arrows indicate cellular elongation observed in both *control* and *Atf3*^*KO*^ cells after one day of IL-4 stimulation (F, G, L, M). While *Atf3*^*KO*^ remain elongated (N, O), *control* cells round up and MGCs (arrowheads) are readily observed in 3-day IL-4 treated culture (H, I). **(P)** Number of nuclei per MGC increases in *control* BMDM cultures over time, following IL-4 stimulation. **(Q)** Plot showing fusion index (fused nuclei/total number of nuclei in %) in *control* and *Atf3*^*KO*^ cells, at one and three days of IL-4 stimulation. *Control* cells form MGCs after one day of IL-4 treatment, and their number increases with time, while *Atf3*^*KO*^ BMDMs do not multinucleate. **(R-U)** Representative bright field and confocal images of BMDMs seeded at higher density and concentration of IL-4 for three days (R, T) or on fusogenic plastic (Permanox) (S, U). While *control* cells form MGCs (R, S), *Atf3*^*KO*^ do not multinucleate in either of the conditions (T, U). **(V-X)** Representative confocal micrographs of peritoneal macrophages stimulated with IL-4 for three days (V, W) and quantification of the fusion index (X) show lack of MGCs in *Atf3*^*KO*^ culture (W) compared to *control* (V). **(Y-EE)** Representative bright field and confocal micrographs show MGCs in *control* and *Atf3*^*KO*^ BMDM cultures stimulated with RANKL for nine days (Y, Z), FSL-1 for seven days (BB, DD) or infected with *Mycobacterium tuberculosis* (Erdman-GFP, cyan) for seven days (CC, EE). RANKL-stimulated *control* and *Atf3*^*KO*^ BMDMs form multinucleated osteoclasts that are positive for TRAP staining (Y, Z). *Atf3*^*KO*^ osteoclasts are smaller in size compared to the *control* (AA). Data information: Data represent means ± s.d.; **p<0.01, ***p<0.001, ****p<0.0001, n.s. = non-significant. Statistical significance was determined using Mann-Whitney test. Each dot represents average value for one mouse (A, Q, X, n ≥ 4) or individual cells from three independent experiments (P, AA). Confocal micrographs are z-projections of multiple confocal sections. Phalloidin labels F-actin; DAPI labels nuclei. Scale bars: 100 µm (D, F, H, J, L, N, R, T, Y, Z) and 20 µm (E, G, I, K, M, O, S-W, BB-EE).

Despite this defect, *Atf3*^*KO*^ BMDMs differentiated into multinucleated osteoclasts in response to RANKL *in vitro*, albeit they were smaller compared to the *control* (Fig. 1Y-AA). *Atf3*-deficient BMDMs also formed MGCs when stimulated with FSL-1 or infected with a wild-type strain of *Mycobacterium tuberculosis* (Erdman-GFP) (Fig. 1BB-EE), demonstrating that the role Atf3 in multinucleation is specific to IL-4 signaling rather than a general requirement for MGC formation. These findings identify IL-4 as a potent inducer of Atf3 and establish Atf3 requirement for IL-4-mediated MGC formation but dispensable for RANKL- or TLR2-driven multinucleation.

### Atf3 promotes macrophage cell-cell fusion in response to IL-4

Given the multinucleation defect of *Atf3*^*KO*^ BMDMs in response to IL-4, we examined whether Atf3 is required for cell-cell fusion or acytokinetic division, the two principal mechanisms generating MGCs (Fig. 2A). Time-lapse imaging revealed that both processes occurred in *control* BMDM cultures upon IL-4 stimulation (Fig. 2A, B and Supplementary Movie 1). To assess their relative contributions, we co-cultured differentially labeled BMDMs expressing either cytoplasmic tdTOMATO or nuclear H2B-EGFP (Fig. 2A). After three days of IL-4, most *control* MGCs (95%) were positive for both fluorescent markers (Fig. 2C, E, F), indicating that cell-cell fusion is the dominant mechanisms underlying IL-4-induced multinucleation. GFP-positive nuclei accounted for nearly 50% of the total nuclei in tdTOMATO-positive MGCs, suggesting equal fusion competence of the two *control* populations (Fig. 2G). In contrast, fusion efficiency was dramatically reduced when *control* BMDMs were co-cultured with *Atf3*^*KO*^ cells (Fig. 2D-F). *Atf3*^*KO*^ cells contributed less than 15% of nuclei to MGCs (Fig. 2G) and co-cultures of *Atf3*^*KO*^ BMDMs alone failed to generate any MGCs (Fig. 2I). The proportion of binucleated cells was also decreased in *Atf3*^*KO*^ cultures, and those that did form never expressed both fluorescent markers (Fig. 2I, J) consistent with incomplete mitosis rather than fusion. Together, these results establish Atf3 as a key regulator of macrophage fusion and reinforce the concept that cell-cell fusion, rather than acytokinetic division, is the primary mechanism driving IL-4-mediated MGC formation.

**Figure 2.**
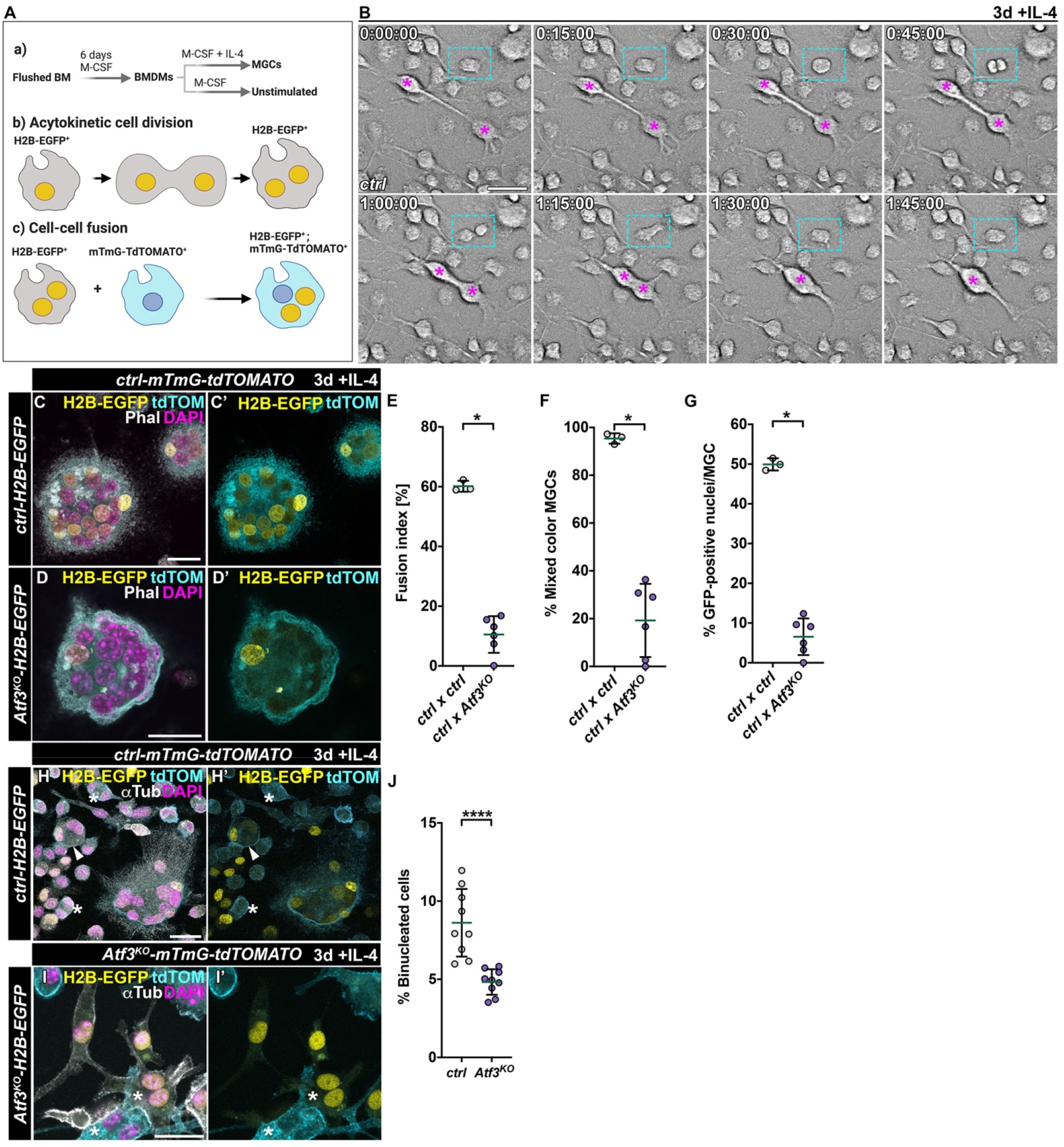
Atf3 is indispensable for IL-4-induced macrophage fusion. **(A)** Schematic representation of the IL-4 stimulation protocol to generate MGCs (a). Cells expressing nuclear H2B-EGFP (yellow) were mixed with cells expressing membrane-targeted mTmG-TdTOMATO (cyan). Both acytokinetic cell division (b) and cell-cell fusion (c) drive multinucleation. **(B)** Bright field single frames from time-lapse live imaging of *control* BMDMs stimulated with IL-4 for three days. Both cell-cell fusion (asterisks) and incomplete cytokinesis (rectangle) drive multinucleation. **(C-G)** Representative confocal micrographs (C, D) and quantifications (E-G) of MGCs formed in co-culture of mTmG-TdTOMATO-expressing *control* cells with *control* (C) or *Atf3*^*KO*^ (d) cells expressing nuclear H2B-EGFP stimulated with IL-4 for three days. Co-culture of *Atf3*^*KO*^ with *control* cells significantly decreases fusion index (E). The percentage of MGCs that express both nuclear H2B-GFP and membrane-targeted mTmG-TdTOMATO is significantly decreased in *ctrl x Atf3*^*KO*^ co-cultures compared to *ctrl x ctrl* ones (F). While *ctrl x ctrl* mixed cultures display TdTOMATO-positive MGCs that enclose 50% H2B-EGFP-positive nuclei in average, the contribution of H2B-GFP-positive *Atf3*^*KO*^ nuclei is below 10%. Plot depicts the contribution of each tagged population to MGCs, as a percentage of GFP-positive nuclei per MGC (G). **(H-J)** Representative confocal micrographs of *control* (H) and *Atf3* (I) co-cultures in which BMDMs expressing nuclear H2B-EGFP were mixed with ones expressing membrane-targeted mTmG-TdTOMATO from the same genotype and stimulated for three days with IL-4. Compared to mTmG-TdTOMATO-positive MGCs containing H2B-EGFP-positive nuclei in *control* co-cultures (H), *Atf3*^*KO*^ BMDMs can only binucleate and express either H2B-GFP or mTmG-TdTOMATO only (asterisk) compared to single (asterisk) and mixed color (arrowdead) binucleated cells in *control* co-cultures (H). The percentage of binucleated cells in *Atf3*^*KO*^ cultures is significantly decreased compared to *control* cultures stimulated with IL-4 for three days (J). Data information: Data represent means ± s.d.; *p<0.05, ****p<0.0001. Statistical significance was determined using Mann-Whitney test (E-G, n ≥ 3; J, n = 9). Each dot represents average value for one mouse. Confocal micrographs are z-projections of multiple confocal sections. Phalloidin labels F-actin, αTubulin labels microtubules, DAPI labels nuclei. Scale bars: 100 µm (B) and 20 µm (C, D, H, I).

### Atf3 acts a critical transcriptional coordinator of IL-4-driven macrophage program

To investigate the underlying cause of the multinucleation defect, we assessed whether *Atf3*^*KO*^ BMDMs could sense and respond to IL-4. Our CRISPR/Cas9-generated homozygous *Atf3*^*KO*^ mice are viable, fertile and phenotypically normal consistent with previous reports (Hartman *et al*, 2004). However, in line with the established anti-inflammatory function of Atf3 (Gilchrist *et al*., 2006), naive *Atf3*^*KO*^ macrophages displayed a heightened basal inflammatory state, characterized by elevated levels of phosphorylated STAT1 (p-STAT1) and increased expression of NF-κB-dependent inflammatory genes, including *Il6* and *Isg15*, compared with *controls*, as shown by immunoblotting (Fig. 3A, B) and RT-qPCR (Fig. 3C). Despite this pro-inflammatory priming, the IL-4 signaling remained functional in *Atf3*-deficient cells. Upon IL-4 stimulation, p-STAT1 levels decreased comparably in both genotypes. Importantly, levels of phosphorylated STAT6 (p-STAT6), the essential transcription factor for IL-4-driven gene expression and fusion (Moreno *et al*, 2007), increased to a similar extent in both *control* and *Atf3*^*KO*^ macrophages (Fig. 3D, E). These data demonstrate that while Atf3 is required to restrain basal inflammation, its absence does not disrupt the primary IL-4 signaling, indicating that the observed fusion defect occurs downstream, parallel to, or independent of STAT6.

**Figure 3.**
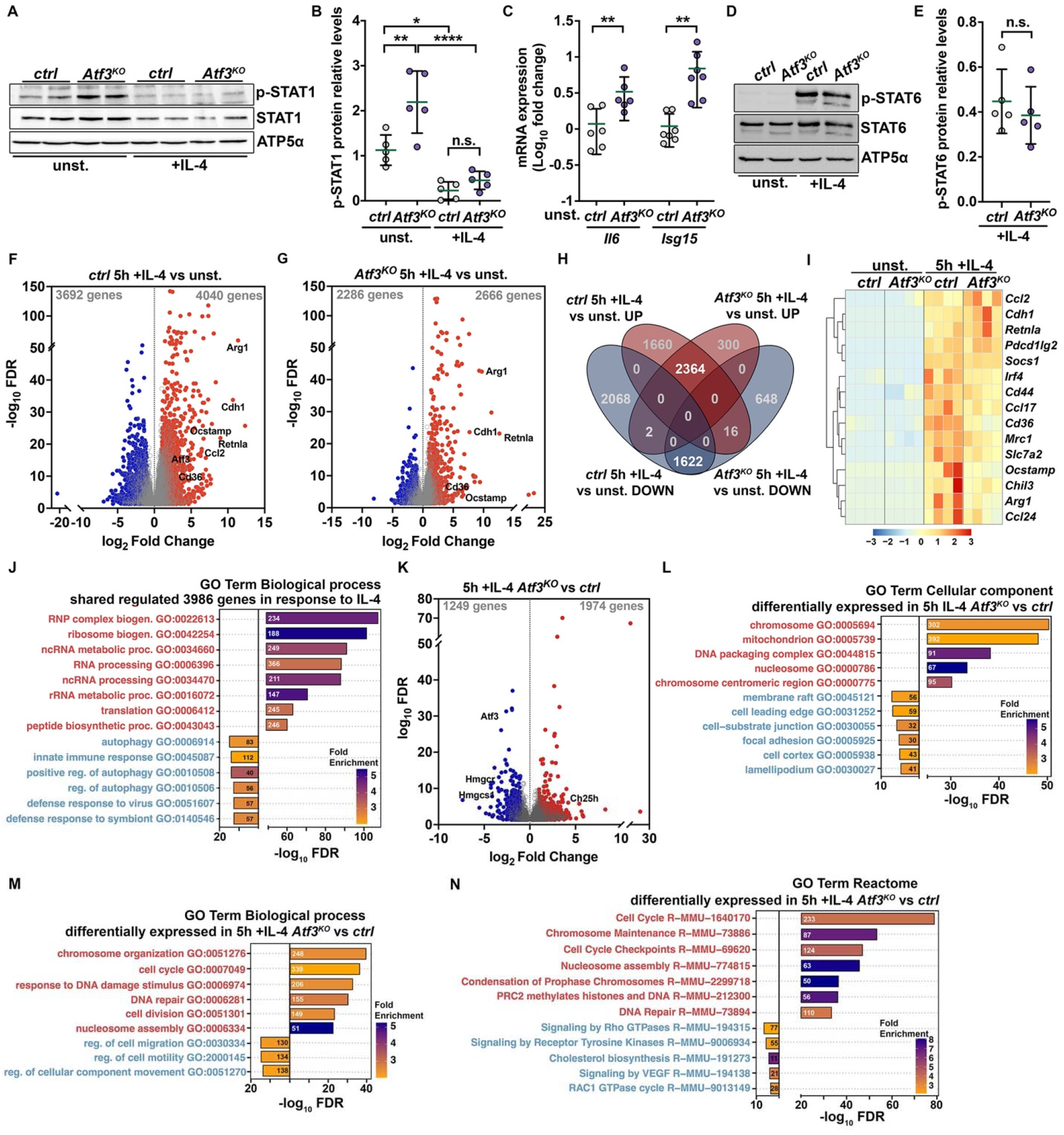
Atf3 deficient BMDMs activate STAT6-mediated transcription as well as unique gene expression signature in response to IL-4. **(A, B)** Representative western blot (A) and quantification (B) showing phosphorylated (pSTAT1) and total STAT1 protein in *control* and *Atf3*^*KO*^ BMDMs in the unstimulated state (unst.) and one day after IL-4 stimulation (+IL-4). The higher pSTAT1 levels in unstimulated *Atf3*^*KO*^ cells decrease upon IL-4 stimulation to levels observed in IL-4 treated *control* BMDMs. ATP5α was used as a loading control. **(C)** RT-qPCR shows increased expression of *Il6* and *Isg15* transcripts in unstimulated *Atf3*^*KO*^ BMDMs compared to *control*. Data are Log_10_ fold changes in expression relative to *control* (ΔΔCt method). Levels of *Gapdh* transcripts were used for normalization. **(D, E)** Representative western blot and quantification showing that both *ctrl* and *Atf3*^*KO*^ BMDMs activate STAT6 (pSTAT6) in response to IL-4. ATP5α was used as a loading control. **(F, G)** Volcano plots depict significantly regulated genes (*pAdj* < 0.05) in *control* (*ctrl*) (F) and *Atf3*^*KO*^ (G) BMDMs stimulated with IL-4 for 5h, compared to the unstimulated state, as detected by RNA sequencing. The total number of downregulated and upregulated genes is indicated in gray for each plot. Dark blue/red colors highlight genes regulated ≥ 2-fold. M2 genes, such as *Arginase-1 (Arg-1), Resistin-like alpha (Retnla)* and cell-cell fusogens, such as *Cadherin (Cdh1), Osteoclast stimulatory transmembrane protein (OC-STAMP)* and *CD36*, are among the most upregulated genes in response to IL-4 in both genotypes. **(H)** Venn diagram shows overlap of upregulated (red) and downregulated (blue) genes in *control* and *Atf3*^*KO*^ BMDMs upon 5h of IL-4 stimulation in comparison to unstimulated state. **(I)** Expression of IL-4-regulated genes, including M2-like and fusion-promoting genes, is induced upon 5h of IL-4 stimulation in comparison to unstimulated cells, in both *control* and *Atf3*^*KO*^ BMDMs. The heatmap represents the Z-scores of normalized RNA-seq counts, scaled by rows. **(J)** Gene ontology (GO) “Biological processes” enrichment plot of the 3986 genes that are significantly regulated (*pAdj* < 0.05) in the same direction (1622 downregulated, blue, and 1622 upregulated red) in IL-4 stimulated *control* and *Atf3*^*KO*^ cells, represented in H. The Fold enrichment for each GO term is color-coded and the number of genes indicated. **(K)** Volcano plot depicts significantly regulated genes (*pAdj* < 0.05) in *Atf3*^*KO*^ BMDMs compared to *controls*, both stimulated with IL-4 for 5h, as determined by RNA sequencing. The total number of downregulated and upregulated genes is indicated in gray with dark blue and red colors highlighting genes regulated ≥ 2-fold. **(L-N)** Cellular component (L), Biological process (M) and Reactome (N) GO term enrichment analysis of differentially regulated genes (*pAdj* < 0.05) in *Atf3*^*KO*^ BMDMs compared to control cells after 5 hours of IL-4 stimulation. The GO terms derived from 1249 significantly downregulated, and 1974 upregulated genes are represented to the left (blue) and to the right (red) of the Y axis, respectively. The Fold enrichment for each GO term is color-coded and the number of genes is indicated. Data information: Data represent means ± s.d.; *p<0.05, **p<0.01, ****p<0.0001, n.s. = non-significant (B, C, E). Statistical significance was determined using ordinary one-way ANOVA with Tukey’s multiple comparison test (B, n = 5) or unpaired t-test assuming unequal variance (Mann-Whitney test) (C, n ≥ 6; E, n = 5). Each dot represents average value for one mouse.

Genome-wide RNA-sequencing on *control* and *Atf3*^*KO*^ BMDMs before and after 5 hours of IL-4 stimulation revealed that IL-4 altered the expression of 7,732 genes (*pAdj* < 0.05) in *control* cells (Fig. 3F and Supplementary Dataset 1), of which 3,986 were regulated similarly in *Atf3*^*KO*^ BMDMs (Fig. 3F-I). The 2,364 commonly upregulated genes were enriched for functional categories related to ribonucleoprotein complex assembly, ribosome biogenesis, RNA processing and translation (Fig. 3J and Supplementary Dataset 2). Consistent with STAT6 activation, this set included canonical targets such as *Arginase-1 (Arg-1), Resistin-like alpha (Retnla)* and *Chitinase-like protein 3 (Chil3)*, as well as fusion-associated genes like *Cadherin (Cdh1), CD44, CD36* and *Osteoclast-stimulatory transmembrane protein (Ocstamp)* (Fig. 3I). Both genotypes shared 1,622 commonly downregulated genes that were enriched for functions associated with autophagy, immune responses, and interferon signaling pathways (Fig. 3J and Supplementary Dataset 2). These findings demonstrate that IL-4 is sufficient to shift *Atf3*^*KO*^ BMDMs from a STAT1-driven pro-inflammatory profile to the canonical STAT6-mediated transcriptional program, including the induction of known fusion-promoting genes.

Despite this overlap and a clear STAT6-regulated signature (Fig. 3H-J), IL-4-treated *Atf3*^*KO*^ BMDMs showed 3,223 differentially expressed genes (DEGs) (*pAdj* < 0.05) compared with IL-4 stimulated *controls*, comprising 1,974 upregulated and 1,249 downregulated transcripts (Fig. 3K). The upregulated genes were overrepresented for nuclear-localized proteins involved in chromatin organization, cell cycle regulation and DNA repair (Fig. 3L-N and Supplementary Dataset 2). Downregulated genes encoded proteins at the cell cortex and plasma membrane, including components of cell junctions, projections (e.g., lamellipodia and leading edges), and membrane microdomains (Fig. 3L). Gene ontology analysis linked these changes to pathways and processes associated with cell motility, receptor tyrosine kinase and small GTPase signaling, and cholesterol metabolism (Fig. 3M, N and Supplementary Dataset 2). Collectively, these results identify Atf3 as a critical transcriptional coordinator of IL-4-driven macrophage program downstream of, or in parallel to STAT6, integrating cytoskeletal architecture, lipid metabolism and genome integrity. While IL-4 effectively rewires the inflammatory state and activates a robust STAT6-driven transcriptional program in *Atf3*^*KO*^ macrophages, dysregulation of Atf3-dependent modules likely represents the mechanistic basis for the persistent fusion defects.

### Atf3 loss impairs cytoskeletal organization and dynamics in IL-4 stimulated macrophages

The marked transcriptional downregulation of structural and regulatory cytoskeletal components in *Atf3*^*KO*^ macrophages (Fig. 3L-N and Supplementary Datasets 1, 2), together with prior evidence of Atf3’s role in cytoarchitecture regulation (Boespflug *et al*., 2014; Donohoe *et al*., 2018; Du *et al*., 2022; Sekyrova *et al*., 2010), prompted us to examine how loss of *Atf3* affects cytoskeletal organization and membrane dynamics, key processes for efficient macrophage fusion.

Following IL-4 stimulation, both *control* and *Atf3*^*KO*^ BMDMs initially adopted the characteristic elongated morphology (Fig. 1F, G, L, M). Over time, *control* BMDMs rounded, extended lamellipodia, and fused (Figs. 1H, I, and 4A), whereas *Atf3*^*KO*^ cells maintained their elongated shape and formed multilayered aggregates with intact intercellular boundaries (Figs. 1N, O, and 4B). In *control* macrophages, IL-4 induced extensive membrane ruffling, lamellipodia and filopodia formation, supported by a dense cortical actomyosin network (Fig. 4C, E, G-H, K and Supplementary Movie 2). In *Atf3*^*KO*^ BMDMs, membranes were smoother with sparse filopodia and actomyosin cables concentrated along stellate extensions (Fig. 4D, F, I-L and Supplementary Movie 3). Biochemical fractionation showed a reduced F/G actin ratio in IL-4-stimulated *Atf3*^*KO*^ BMDMs relative to *control* cells (Fig. 4P and Supplementary Fig. 1A). These changes were accompanied by alteration in actin regulators, including elevated levels of the Wiskott–Aldrich syndrome protein (WASP), increased active (dephosphorylated) Cofilin, and abnormal perinuclear accumulation of the actin-crosslinking protein Filamin A (FLNA) (Fig. 4M-O, Q-T and Supplementary Fig. 1B).

**Figure 4.**
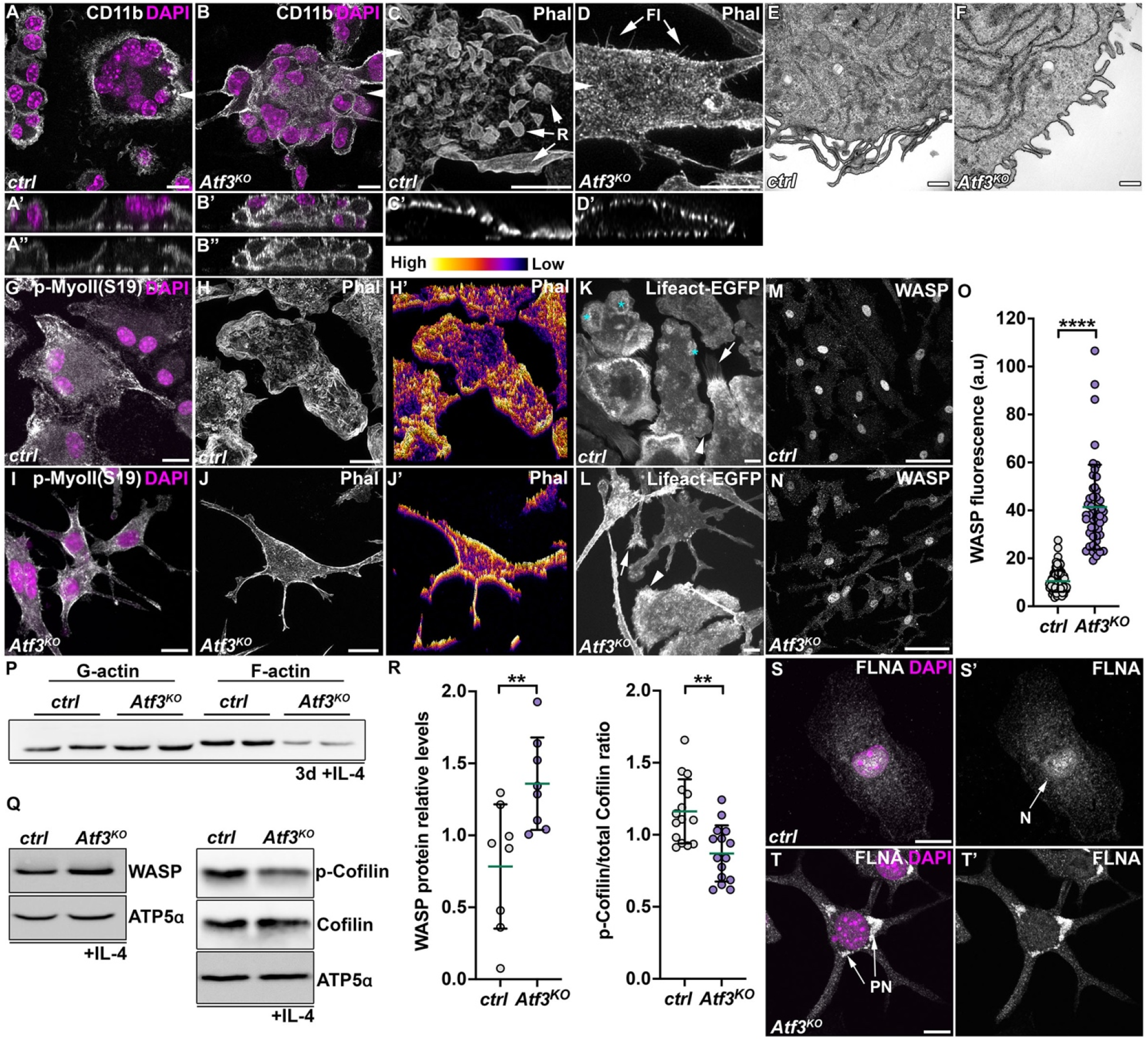
Atf3 loss affects cytoskeletal responses of macrophages to IL-4. **(A-B)** Representative confocal micrographs of IL-4 stimulated BMDMs stained against the membrane-localized integrin CD11b show numerous DAPI-stained nuclei within a single *control* BMDM (A-A’’). In contrast, *Atf3*^*KO*^ BMDMs form multilayered aggregates, with mononuclear cells clearly separated by cell membranes (B-B’’). White arrowheads indicate the positions of the cross-sections. **(C-D)** 3D confocal projection of z-stack images of *control* (C) BMDMs stained for F-actin (Phal) after three days of IL-4 stimulation shows prominent ruffling (R), while *Atf3*^*KO*^ BMDMs display scarce thin filopodia (Fl, arrow). White arrowheads indicate the positions of the cross-sections, which are presented below (C’, D’). **(E-F)** Electron microscopy images highlight long filopodia in *control* BMDMs treated with IL-4 for three days (e), compared to the shorter projections in *Atf3*^*KO*^ cells (F). **(G-J)** Compared to the rich cortical actomyosin network in *control* cells, visualized by staining for F-actin (Phal) (G, H, corresponding intensity heatmaps G’, H’) and phospho-Myosin (S19), actomyosin cables in *Atf3*^*KO*^ BMDMs (I, J) are concentrated beneath the plasma membrane and extend along the branches. **(K-L)** Confocal micrographs of BMDMs expressing Lifeact-EGFP reveal actin-rich podosome rings (K, asterisks), lamellipodia (K, arrowhead) and filopodia (K, arrow) in *control* cells while these structures are notably reduced in *Atf3*^*KO*^ BMDMs (l). **(M-O)** Representative confocal micrographs (M, N) and quantification (O) of IL-4 stimulated BMDMs showing increased WASP fluorescence intensity in *Atf3*^*KO*^ macrophages (N) compared with *control* cells (M). In contrast to the primarily nuclear signal in *control* BMDMs, *Atf3*^*KO*^ BMDMs display WASP signal in both nucleus and cytoplasm. **(P)** Representative western blot of IL-4 treated cells shows marked decrease of F-actin in *Atf3*^*KO*^ BMDMs compared to *control*. **(Q-R)** Representative western blots of WASP, phospho-Cofilin (p-Cofilin) and total Cofilin protein levels (Q) and quantification (R) in IL-4-stimulated BMDMs of the indicated genotypes. ATP5α was used as a loading control. **(S-T)** Representative confocal micrographs reveal differential localization and enrichment of Filamin A (FLNA) in the nucleus (arrow, N) of *control* (S, S’) cells and in perinuclear foci (PN) of *Atf3*^*KO*^ (T, T’) BMDMs stimulated by IL-4. Data information: Data represent means ± s.d.; **p<0.01, ****p<0.0001 (O, R). Statistical significance was determined using Mann-Whitney test. Each dot represents individual cell from three independent experiments (O, n = 3) or individual animal (R, n = 8 left, n = 15 right) Micrographs are z-projections of multiple confocal sections. DAPI labels nuclei. Scale bars: 10 µm (A-D, G-L, S, T), 50 µm (M, N), and 0.5 µm (E, F).

These results implicate Atf3 as a critical regulator of actin cytoskeleton organization and turnover downstream of IL-4 signaling and suggest that defective cytoskeleton remodeling underlies the fusion defects in *Atf3*^*KO*^ macrophages.

### Atf3 controls the mevalonate pathway required for protein prenylation and cholesterol homeostasis

Actin regulators, such as WASP, Cofilin and FLNA are controlled by the Rho family GTPases (RhoA, Rac1 and Cdc42) (Jaffe & Hall, 2005). The membrane localization and activity of these GTPases require post-translational farnesylation or geranylgeranylation by prenyltransferases (Hodge & Ridley, 2016). The required isoprenoids are synthesized via the mevalonate pathway from farnesyl pyrophosphate (FPP), a common precursor for non-sterol isoprenoids and cholesterol (Fig. 5A and Supplementary Fig. 1C). Inhibition of prenyltransferases, or the rate-limiting enzyme of the mevalonate pathway, HMG-CoA reductase (Hmgcr), disrupts Rho GTPase membrane association and activity resulting in altered actin turnover and remodeling (Akula *et al*, 2019; Cordle *et al*, 2005).

**Figure 5.**
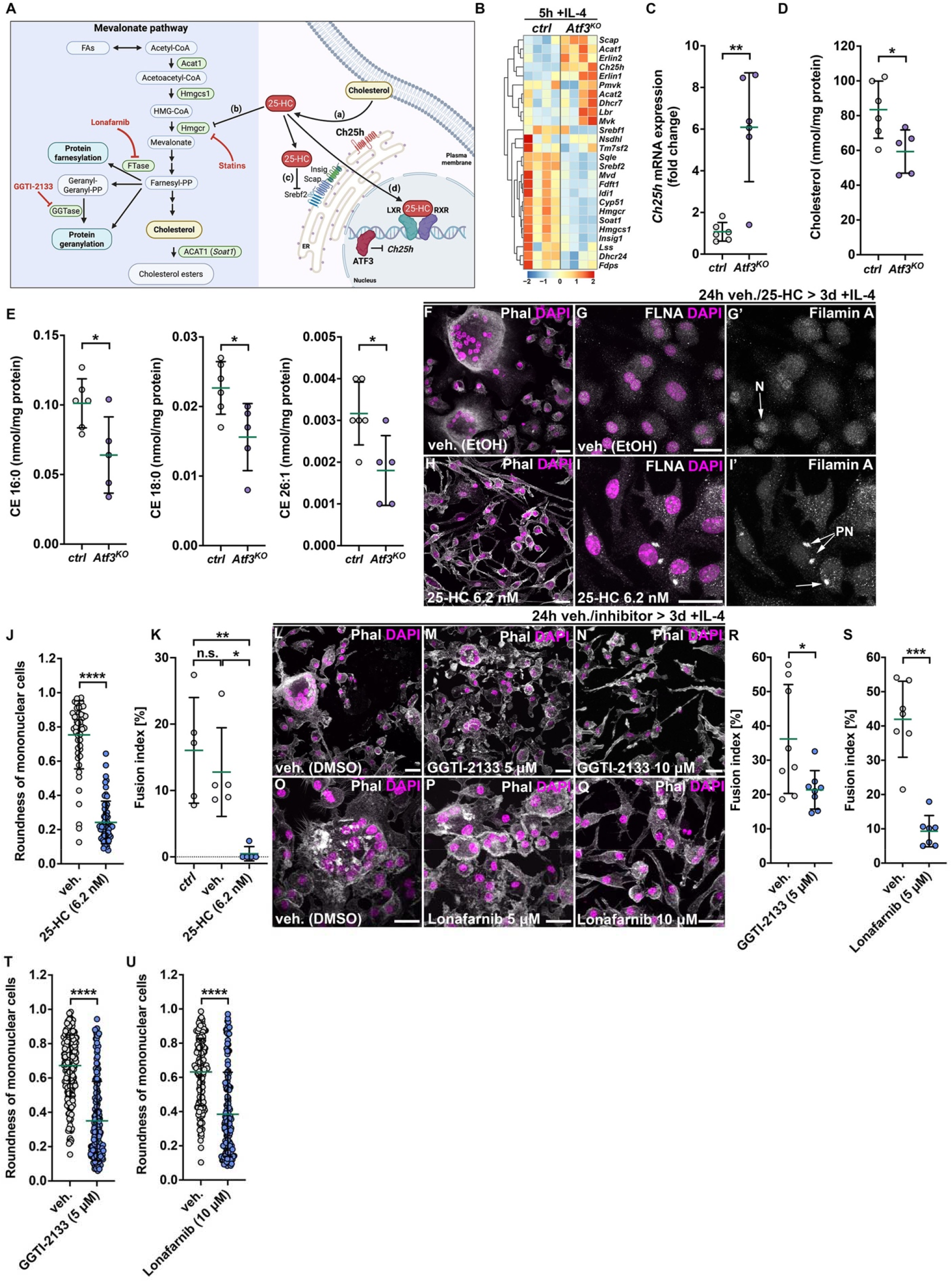
25-HC-mediated inhibition of the mevalonate pathway and protein prenylation recapitulates the phenotypes of Atf3 loss. **(A)** Schematic representation of cholesterol and isoprenoid metabolism in the cell, highlighting the mevalonate pathway. The ER-resident enzyme Cholesterol-25-hydroxylase (Ch25h) generates the oxysterol 25-Hydroxycholesterol (25-HC). 25-HC causes sequestration of a free cholesterol pool from the plasma membrane (a). 25-HC regulates cholesterol biosynthesis by suppressing the rate-limiting enzyme of the mevalonate pathway Hmgcr (b) and by binding the Insulin-induced gene (INSIG) and forming a complex that retains Sreb2-Scap inactive in the ER (c). In addition, 25-HC binds LXR-RXR transcription factor dimers to initiate expression of genes involved in the absorption, degradation, transportation, and excretion of cholesterol (d). Besides *de novo* synthesis of cholesterol, the mevalonate pathway also generates isoprenoids. Isoprenoid intermediates, farnesyl pyrophosphate and geranylgeranyl pyrophosphate, are used for prenylation of various proteins. Compounds inhibiting the mevalonate pathway (25-HC, Statins) and protein prenylation (GGTI-2133, Lonafarnib) are depicted. The model was generated using BioRender.com. **(B)** Heatmap shows differential expression of selected genes associated with cholesterol biosynthesis in *control* and *Atf3*^*KO*^ BMDMs stimulated with IL-4 for 5h. The heatmap represents the Z-scores of normalized RNA-seq counts, scaled by rows. **(C)** RT-qPCR shows upregulation of *Ch25h* expression in IL-4 stimulated *Atf3*^*KO*^ BMDMs compared to *control*. **(D, E)** Total cholesterol levels, quantified by LC-MS/MS (D), and the three main classes of cholesterol ester species, CE 16:0, CE 18:0 and CE 26:1 (E), are significantly decreased in IL-4 stimulated *Atf3*^*KO*^ BMDMs compared to *control*. Each dot represents one mouse. Measurements from independent experiments were pooled and normalized to average per experiment. **(F-K)** Representative confocal micrographs (F-I) and quantifications (J, K) showing 3-day IL-4-stimulated *control* BMDMs pre-treated with EtOH (vehicle, veh.) (F, G) or 25-HC (6.2 nM) (H, I) for 24h. In contrast to MGC formation (F, K) and nuclear localization of Filamin A (G’) in vehicle treated cells, cells exposed to 25-HC remain mononucleated (H, K), elongate (H, J) and Filamin forms perinuclear foci (I’). **(L-U)** Representative micrographs (L-Q) and quantifications of cell fusion (R, S) and cell roundness (T, U) of *control* BMDMs pre-treated for 24 with DMSO (vehicle, veh.) (L, O), 5 µM (M, P), or 10 µM (N, Q) of an inhibitor of geranylgeranylation, GGTI-2133 (M, N), or farnesylation, Lonafarnib (P, Q), prior to IL-4 stimulation. Treatment with GTI-2133 and Lonafarnib reduces MGC formation (R, S) and cells are more elongated (T, U) compared to vehicle treated BMDMs. Data information: Data represent means ± s.d.; *p<0.05, **p<0.01, ***p<0.001, ****p<0.0001, n.s. = non-significant. Statistical significance was determined using unpaired t-test assuming unequal variance (Welch’s test) (C-E) and Mann-Whitney test (J, K, R-U). Each dot represents average value of one mouse (C-E, K, R, S, n ≥ 5) or single cells from 3 independent experiments (J, T, U). Micrographs are z-projections of multiple confocal sections. Phalloidin labels F-actin; DAPI labels nuclei. Scale bars: 20 µm.

*Atf3*^*KO*^ macrophages showed downregulation of genes associated with small GTPase regulation and signaling (Fig. 3N and Supplementary Dataset 2), and specifically the mevalonate pathway, including *Hmgcs1* and *Hmgcr*, the cholesterol biosynthesis regulator, Sterol Regulatory Element Binding Protein 2 (*Srebf2*), and a Sterol O-acyltransferase 1 (*Soat1*), which esterifies cholesterol (Figs. 3N, 5B, Supplementary Fig. 1C and Supplementary Dataset 2). In contrast, *Ch25h*, a known direct Atf3 target gene encoding Cholesterol-25-hydroxylase (Gold *et al*., 2012), was strongly upregulated in *Atf3*^*KO*^ BMDMs compared to *control* cells (Figs. 3K, 5B, 5C, and Supplementary Dataset 1). Its oxysterol product, 25-hydroxycholesterol (25-HC), depletes accessible plasma membrane cholesterol, promotes LXR-mediated efflux, suppresses Srebf2 activation, and accelerates Hmgcr degradation (Fig. 5A and Supplementary Figure 1C). Consistent with these molecular changes, total cholesterol and major cholesteryl esters (CE 16:0, CE 18:0 and CE26:1) were significantly reduced in IL-4 stimulated *Atf3*^*KO*^ BMDMs compared to *controls* (Fig. 5D, E). These findings identify Atf3 as a central regulator of the mevalonate pathway in IL-4-stimulated macrophages, modulating isoprenoid and cholesterol biosynthesis essential for the cytoskeletal and membrane remodeling underlying macrophage fusion. The strong upregulation of *Ch25h* in *Atf3*-deficient macrophages points to its oxysterol product, 25-HC, as a potential mediator of these effects.

### Elevated 25-HC impairs MGC formation and phenocopies morphological changes caused by *Atf3*-deficiency

To determine whether elevated 25-HC can replicate the cytoskeletal and fusion defects of *Atf3*-deficient macrophages, we pre-treated control BMDMs with 25-HC for 24 hours prior to IL-4-stimulation. Remarkably, 25-HC induced elongated, stellate morphology and perinuclear FLNA foci (Fig. 5F-J), closely resembling the phenotypes of *Atf3*^*KO*^ BMDMs following IL-4 stimulation (Figs. 1O, 4I-L, 4T). Notably, 25-HC treatment suppressed MGC formation (Fig. 5F, H, K). Moreover, similar morphological changes and reduced multinucleation were also observed when *control* BMDMs were treated with inhibitors of geranylgeranylation (GGTI-2133) and farnesylation (Lonafarnib) (Fig. 5A, 5L-U).

These results implicate the upregulation of Ch25h, and accumulation of 25-HC driven by Atf3 deficiency in the regulation of macrophage morphology, cytoskeletal remodeling and fusion capacity. The similarity of phenotypes elicited by prenylation inhibitors further supports a model in which 25-HC suppresses the mevalonate pathway, limiting isoprenoid availability and thereby impairing Rho GTPase-dependent actin remodeling required for IL-4-induced multinucleation.

### Blocking Ch25h in *Atf3*^*KO*^ BMDMs rescues cell morphology but not cell fusion

The strong phenotypic similarity between 25-HC-treated control macrophages and *Atf3*-deficient cells suggested that excess 25-HC is a major driver of the cytoskeletal abnormalities in *Atf3*^*KO*^ BMDMs. To directly test this hypothesis, we generated *Ch25h*-deficient mice using CRISPR/Cas9 (Fig. 6A). Homozygous *Ch25h*^*KO*^ mice are born at the expected Mendelian ratio, producing fertile adults without noticeable phenotypes. Unlike IL-4-stimulated *Atf3*^*KO*^ BMDMs, which had reduced cholesterol, *Ch25h*^*KO*^ macrophages maintained cholesterol levels comparable to *controls* (Fig. 6B) despite a marked increase in Hmgcr protein (Fig. 6C, D). Unexpectedly, *Ch25h*^*KO*^ macrophages showed elevated *Atf3* protein levels in both basal and IL-4 stimulated conditions (Fig. 6C, E) and enhanced fusion capacity, forming more and larger MGCs (Fig. 6F-J). These observations suggest a feedback loop between Atf3 and Ch25h, in which 25-HC limits fusion competency.

**Figure 6.**
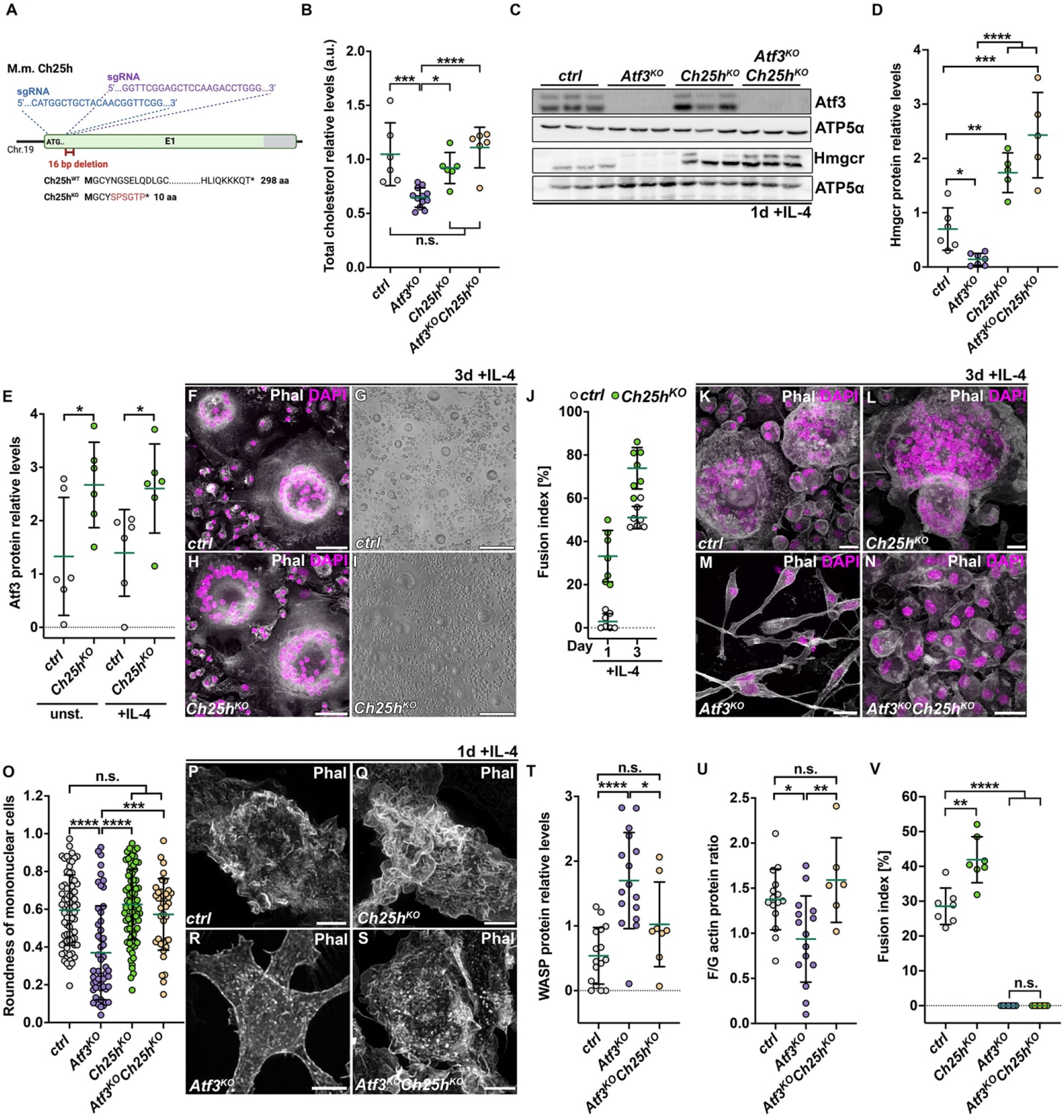
Loss of Ch25h rescues cell morphology and F-actin levels in Atf3 deficient BMDMs. **(A)** Schematic representation of the *Ch25h* locus targeted by two sgRNAs to generate a 49 bp deletion resulting in a frameshift and the introduction of a premature STOP codon. Box represents a single exon 1 (E1) with a start codon (ATG) and 3’UTR sequence (grey). **(B)** Decreased total cholesterol levels in *Atf3*^*KO*^ BMDMs are normalized to *control* levels upon simultaneous loss of Ch25h. Loss of Ch25h alone does not alter cholesterol levels. Measurements from independent experiments were pooled and normalized to the mean per experiment. **(C-E)** Representative western blots (C) and quantification (D, E) show elevated Hmgcr protein in IL-4 stimulated *Ch25h*^*KO*^ and *Atf3*^*KO*^*Ch25h*^*KO*^ BMDMs while reduced in *Atf3*^*KO*^ cells (D). Atf3 protein is increased in 1-day IL-4 stimulated *Ch25h*^*KO*^ BMDMs relative to *control* (E). ATP5α was used as a loading control. Each genotype is represented by three BMDM samples prepared from independent animals. **(F-J)** Representative confocal (F, H) and bright-field (G, I) micrographs of MGCs and quantification of fusion index (fused nuclei/total nr of nuclei in %) (J) showing enhanced multinucleation in *Ch25h*^*KO*^ BMDMs cultures compared to *control* at one and three days of IL-4 stimulation. **(K-O)** Compared to large MGCs formed in *control* (K) and *Ch25h*^*KO*^ (l) BMDM cultures stimulated for three days with IL-4, MGCs are absent in *Atf3*^*KO*^ (M) and *Atf3*^*KO*^*Ch25h*^*KO*^ (N) samples. In contrast to elongated *Atf3*^*KO*^ cells (M, O), *Atf3*^*KO*^*Ch25h*^*KO*^ (N, O) display round morphology like *control* cells (K, O). **(P-S)** Representative images of Phalloidin-stained (Phal) BMDMs show rich F-actin ruffles in IL-4-stimulated *control* (P), *Ch25h*^*KO*^ (Q) and *Atf3*^*KO*^*Ch25h*^*KO*^ (S) cells contrasting with sparse ruffles in *Atf3*^*KO*^ (R) BMDMs. **(T)** Quantification of WASP protein levels by western blot in 1-day IL-4-stimulated BMDMs shows a significant increase in *Atf3*^*KO*^ cells but not in *Atf3*^*KO*^*Ch25h*^*KO*^ cells compared to *control*. See Supplementary Fig. 2D for a representative western blot. **(U)** Quantification of F/G-actin ratio from western blots reveals restored F-actin levels in IL-4-stimulated *Atf3*^*KO*^*Ch25h*^*KO*^ BMDMs contrasting with the decrease observed in *Atf3*^*KO*^ cells. See Supplementary Fig. 2E for a representative western blot. **(V)** Neither *Atf3*^*KO*^ nor *Atf3*^*KO*^*Ch25h*^*KO*^ BMDMs form MGCs in contrast to the enhanced multinucleation observed in *Ch25h*^*KO*^ BMDMs compared to *control*. The plot shows fusion index (fused nuclei/total nr of nuclei in %) in BMDMs stimulated with IL-4 for three days. Data information: Data represent means ± s.d.; *p<0.05, **p<0.01, ***p<0.001, ****p<0.0001, n.s. = non-significant. Statistical significance was determined using ordinary one-way ANOVA with Holm-Šídák’s multiple comparison test (B, D, O, T-V) or unpaired t-test assuming unequal variance (Welch’s test) (E). The dots represent value from individual animals (B, n ≥ 5; D, n ≥ 5; T, n ≥ 8; V, n = 7), except (O), where single cells from 3 independent experiments are depicted, and (J) showing technical replicates from one of the three independent experiments. Micrographs are z-projections of multiple confocal sections. Phalloidin labels F-actin; DAPI labels nuclei. Scale bars: 200 µm (G, I), 50 µm (F, H), 20 µm (K-N), 5 µm (P-S).

To determine whether *Ch25h* loss could reverse the *Atf3*^*KO*^ phenotypes, we generated *Atf3*^*KO*^*Ch25h*^*KO*^ mice. Double knockout mice, like the single gene knockouts, were viable and exhibited no noticeable defects (Supplementary Fig. 2A). In contrast to *Atf3*^*KO*^ BMDMs, *Atf3*^*KO*^*Ch25h*^*KO*^ macrophages showed increased *Hmgcs1* and *Hmgcr* mRNA and protein levels compared to *controls* after IL-4 stimulation (Fig. 6C, d and Supplementary Fig. 2B, C). Importantly, cholesterol levels in double knockouts were comparable to those in *controls* (Fig. 6B), consistent with the phenotype observed in *Ch25h*^*KO*^ cells. Morphologically, *Atf3*^*KO*^*Ch25h*^*KO*^ BMDMs resembled *control* macrophages, rounding and spreading upon IL-4 stimulation (Fig. 6K-O), and regaining membrane ruffling, though less prominent than in *Ch25h*^*KO*^ or *control* cells (Fig. 6P-S). This partial restoration of membrane dynamics coincided with reduced WASP levels and improved actin turnover, as reflected by recovery of the F/G-actin ratio (Fig. 6T, U and Supplementary Fig. 1D, E). Despite these improvements, *Atf3*^*KO*^*Ch25h*^*KO*^ BMDMs did not form MGCs (Fig. 6V).

The observation that *Ch25h* loss enhanced MGC formation in fusion-competent cells, supports a role for 25-HC as a potent modulator of cell-cell fusion, consistent with its established function in viral-cell membrane fusion. These findings highlight the importance of Atf3-mediated *Ch25h* regulation for IL-4-driven cytoskeletal remodeling through control of the mevalonate pathway, while also demonstrating that additional Atf3-dependent mechanisms, beyond cholesterol and isoprenoid metabolism, are required for macrophage multinucleation.

### Atf3 regulation of nuclear lamina morphology and integrity is Ch25h independent

Although actin cytoskeleton organization and membrane dynamics were restored in *Atf3*^*KO*^*Ch25h*^*KO*^ macrophages, these cells failed to fuse, suggesting that loss of *Atf3* affects other structural systems essential for macrophage function. The cytoplasmic actin network is mechanically and functionally coupled to the nucleoskeleton, which is primarily composed of lamin intermediate filaments. Lamins provide nuclear structural stability, mediate mechanotransduction, and regulate chromatin organization, transcription, and genome integrity (Goldman *et al*, 2002; Gruenbaum & Foisner, 2015; Kalukula *et al*, 2022; Miroshnikova *et al*, 2017). Our previous work identified *Drosophila Lamin C*, the ortholog of mammalian *LMNA*, as a direct transcriptional target of Atf3, establishing Atf3 as a positive regulator of *Lamin C* expression (Donohoe *et al*., 2018). Together with the observed enrichment of DNA damage-associated genes in Atf3-deficient BMDMs (Fig. 3l-n), this prompted us to examine macrophage nuclear structure and its properties.

Compared to the oval-shaped nuclei of *control* BMDMs (Fig. 7A-C), *Atf3*^*KO*^ nuclei displayed pronounced morphological distortions, including deep nuclear envelope invaginations (Fig. 7D-G, I). Increased nuclear deformability was accompanied by a higher proportion of γH2A.X-positive nuclei and micronuclei (Fig. 7D, H, J, K), indicative of compromised genome stability. RT-qPCR and immunoblotting revealed significantly reduced *LMNA* mRNA (Fig. 7L) and lamin A/C protein levels (Fig. 7M, N) in *Atf3*^*KO*^ macrophages relative to *control*. Importantly, lamin A/C remain reduced in *Atf3*^*KO*^*Ch25h*^*KO*^ macrophages (Fig. 7M, N), demonstrating that this defect is independent of Ch25h.

**Figure 7.**
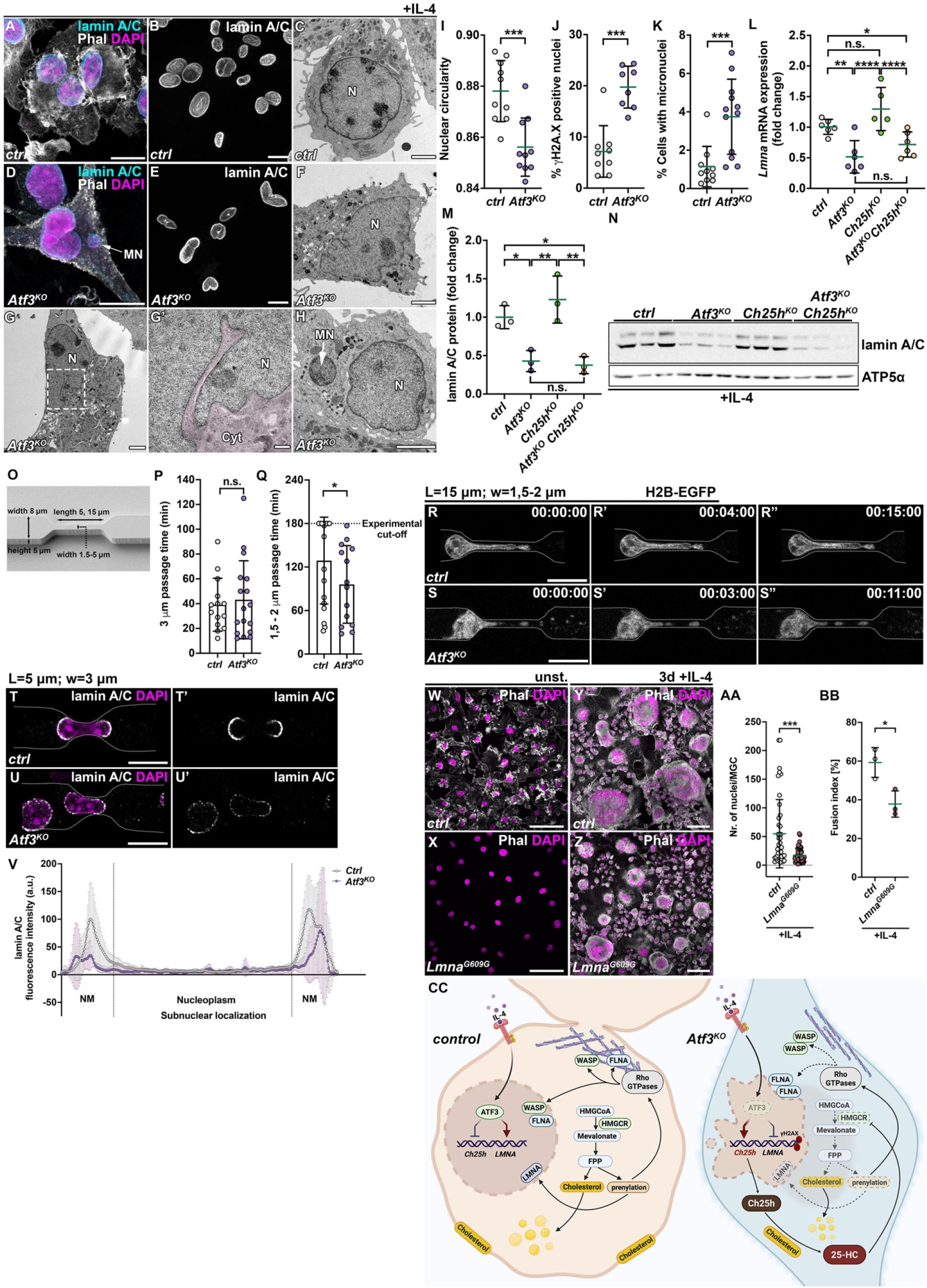
Atf3 controls nuclear morphology and genome integrity of BMDMs independent of Ch25h. **(A-H)** Representative confocal (A, B, D, E) and electron microscopy (C, F, G, H) micrographs of IL-4 stimulated BMDMs highlight differences in nuclear morphology between *control* (A-C) and *Atf3*^*KO*^ cells (D-H), including the presence of micronuclei (MN) (D, H) and nuclear envelope invagination containing cytoplasm in *Atf3*^*KO*^ cells (G, G’). Nuclear lamina is visualized by immunostaining against lamin A/C (A, B, D, E). **(I)** Plot depicting the reduced nuclear circularity of *Atf3*^*KO*^ BMDMs compared to *control* after IL-4 stimulation. Nuclear circularity is defined by the formula 4π(area/perimeter^2^), in which a value of 1.0 indicates a perfect circle. **(J)** Percentage of nuclei positive for the DNA damage marker γH2A.X is increased in *Atf3*^*KO*^ BMDM cultures compared to *control*. **(K)** Percentage of cells containing micronuclei, an extra-nuclear DNA enclosed in lamin A/C-positive membrane, is increased in *Atf3*^*KO*^ BMDM cultures compared to *control*. **(L)** RT-qPCR shows marked downregulation of *Lmna* expression in IL-4 stimulated *Atf3*^*KO*^ BMDMs compared to *control* that is not rescued by simultaneous loss of *Ch25h*. **(M, N)** Quantification (m) and representative western blot (n) show significant reduction of lamin A/C protein in IL-4-stimulated *Atf3*^*KO*^ and *Atf3*^*KO*^*Ch25h*^*KO*^ BMDMs relative to *control*. ATP5α was used as a loading control. **(O)** Schematic representation of 4D-cell microchannels used in migration experiments, with constrictions of variable lengths (L = 5, 15 µm) and widths (w = 1.5 - 5.0 µm). **(P, Q)** The plots display the time (in minutes) required for BMDM nuclei of the depicted genotype to fully traverse constrictions of 3 µm (P) and 1.5-2 µm (Q) in width. *Atf3*^*KO*^ BMDM nuclei show a higher passage rate through the narrower 1.5-2 µm (Q) constrictions compared to *control*. **(R, S)** Sequential images of *control* (R) and *Atf3*^*KO*^ (S) cells expressing H2B-GFP as they pass through 1.5-2 µm constrictions. Nucleo-cytoplasmic GFP leakage is observed in *Atf3*^*KO*^ nuclei over time. **(T-V)** Representative confocal micrographs display lamin A/C distribution in fixed *control* (T, T’) and *Atf3*^*KO*^ (U, U’) BMDM nuclei traversing through 3 µm microchannel constrictions. Plot of LMNA/C intensity across the nucleus (V), as shown in T’ and U’. NM= nuclear membrane. Data points represent quantification of lamin A/C fluorescence intensity from z-stacks. **(W-BB)** Representative confocal micrographs of unstimulated and IL-4 stimulated *control* (W, Y) and *Lmna*^*G608G*^ (X, Z) BMDMs and quantifications (AA, BB) show that *Lmna*^*G608G*^ BMDMs form significantly smaller and fewer MGCs following IL-4 stimulation. **(CC)** Model summarizing the dual checkpoint role of Atf3 in IL-4/STAT6-mediated MGC formation. a) Lipid metabolic control: Atf3 represses Ch25h to limit production of 25-hydroxycholesterol (25-HC), thereby sustaining mevalonate pathway activity, cholesterol availability, and isoprenoid biosynthesis required for protein prenylation including Rho GTPase acting on key actin regulators governing actin cytoskeleton remodeling (WASP, Cofilin, FLNA) and nuclear localization of WASP. b) Nuclear structural integrity: Independently of Ch25h, Atf3 maintains lamin A/C expression, preserving nuclear morphology, mechanical resilience, and genome stability. Loss of Atf3 disrupts both pathways, impairing membrane ruffling, actin dynamics, and nuclear stability, resulting in complete failure of IL-4–driven cell-cell fusion. The model was generated using BioRender.com. Data information: Data represent means ± s.d.; *p<0.05, **p<0.01, ***p<0.001, ****p<0.0001, n.s. = non-significant. Statistical significance was determined using Mann-Whitney (I-K, P, Q) or Welch’s (AA, BB) test assuming unequal variance, or ordinary one-way ANOVA with Tukey’s multiple comparison test (L, M). The dots represent average value from individual animals (I-M, BB, n ≥ 3), except (P, Q, AA), where cells/nuclei from 3 independent experiments are depicted. At least 300 nuclei were counted per individual animal (I-K). Micrographs are z-projections of multiple confocal sections (A, B, D, E, T, U, W-Z, CC, DD). Phalloidin labels F-actin; DAPI labels nuclei. Scale bars: 100 µm (Y, Z), 50 µm (W, X, CC, DD), 10 µm (A, B, D, E, R-U), 2.5 µm (C, F, G, H) and 500 nm (G’).

To determine the functional consequences, we assessed resilience of IL-4 stimulated *control* and *Atf3*^*KO*^ BMDM nuclei during macrophage migration through microfabricated channels of varying widths (Fig. 7O). Tracking nuclear movement via H2B-GFP expression showed comparable transit rates of *control* and *Atf3*^*KO*^ nuclei through wide channels (w ≥ 3 µm) (Fig. 7P). In narrow constrictions (w ≤ 2 µm), all *Atf3*^*KO*^ nuclei, but only around 56% of *control* nuclei, completed passage within a 180-minute experimental window (Fig. 7Q). Despite increased compliance, *Atf3*^*KO*^ nuclei were more fragile, rupturing and leaking H2B-GFP, whereas *controls* primarily exhibited confined nuclear blebbing (Fig. 7R, S, Supplementary Movies 4, 5). Immunostaining revealed lamin A/C enrichment at unconstrained nuclear tips in both genotypes, but overall levels were markedly reduced in protruding *Atf3*^*KO*^ nuclei (Fig. 7T-V).

To directly test whether nuclear lamina defects impair multinucleation, we examined BMDMs from *Lmna*^*G609G*^ knock-in mice, which carry the Hutchinson-Gilford progeria syndrome (HGPS) mutation (Osorio *et al*, 2011). Homozygous *Lmna*^*G609G*^ mice accumulate the truncated prelamin A variant progerin and exhibit reduced levels of mature lamin A/C. Notably, the *Lmna*^*G609G*^ BMDMs showed significantly impaired IL-4-induced MGC formation, producing smaller MGCs compared to *controls* (Fig. 7W-BB).

These results together demonstrate that Atf3 is required for proper *LMNA* expression and nuclear lamina integrity independently of *Ch25h*. We propose that loss of *Atf3*, likely through its effect on Lmna regulation, disrupts the nucleoskeleton, impairing mechanotransduction and compromising genome stability. Together with cytoskeletal abnormalities and altered lipid homeostasis, these defects contribute to the multinucleation failure observed in *Atf3*-deficient macrophages (Fig. 7CC).

## DISCUSSION

Macrophage multinucleation is a striking cellular transformation observed both in physiological tissue remodeling and chronic inflammatory responses (Anderson *et al*., 2008; Brooks *et al*., 2019; Helming & Gordon, 2009; Milde *et al*, 2015; Miron & Bosshardt, 2018; Pereira *et al*, 2018). Here, we identify Atf3 as a pivotal transcriptional regulator of IL-4-driven macrophage multinucleation via cell-cell fusion, a process central to foreign body giant cell (FBGC) formation (Kao *et al*, 1995; McNally & Anderson, 1995). Our findings support a model in which Atf3 promotes fusion through two converging mechanisms: (1) Regulation of lipid metabolism via repression of *Ch25h*, thereby sustaining cholesterol homeostasis and actin remodeling, and (2) Maintenance of nuclear lamina integrity, ensuring nuclear resilience and effective mechanotransduction (Fig. 7W).

### Atf3 as a context-specific regulator of IL-4-driven cell-cell fusion

The requirement for Atf3 in IL-4-induced multinucleation contrasts with its dispensability for RANKL-driven osteoclastogenesis or mycobacteria-induced TLR2-mediated Langhans cell formation. Its strong induction by IL-4, but not by RANKL or TLR2 agonists underscores its specificity for the FBGC program. Mechanistic differences in the formation of distinct MGC subtypes likely contribute to this selectivity. FBGCs and osteoclasts arise primarily via cell-cell fusion (Faust *et al*., 2019; McNally & Anderson, 1995; Stewart *et al*, 2024; Yagi *et al*, 2005), whereas Langhans cells form mainly via modified cell divisions and mitotic defects (Herrtwich *et al*., 2016). This may explain why Atf3-deficient macrophages can still generate MGCs in response to mycobacteria or bacterial lipoproteins. In a Th2-polarized environment favoring FBGC formation, Atf3 functions as a transcriptional node linking IL-4 signaling to fusion-competent cellular programs. Interestingly, the reduced size of Atf3-deficient osteoclasts suggests a modulatory role in the RANKL-induced MGC program, potentially in cooperation with the closely related bZIP transcription factor Jdp2 (Maruyama *et al*, 2012).

The occasional integration of *Atf3*-deficient macrophages into MGCs when co-cultured with control cells indicates that their fusion defect is not absolute. Such residual activity aligns with the model of heterogeneous “founder” and “follower” subpopulations described in myoblast fusion (Kim *et al*, 2015) and later recognized in osteoclasts (Hobolt-Pedersen *et al*, 2014; Levaot *et al*., 2015; Yagi *et al*., 2005), and FBGCs (Faust *et al*., 2019). Founder cells initiate fusion and undergo extensive cytoskeletal remodeling, implying that Atf3 may be essential for establishing the actin- and adhesion-based architecture required for this role.

### Lipid-cytoskeleton coupling in fusion competence

Loss of Atf3 leads to pronounced cytoskeletal abnormalities despite IL-4-STAT-6-mediated transcriptional priming and induction of canonical fusion-associated genes. *Atf3*-deficient macrophages exhibited elongated morphology, reduced membrane ruffling, defective lamellipodia and impaired podosome formation, phenotypes reminiscent of disruptions in Rho GTPase family members (RhoA/RhoB, Cdc42 and Rac1) or the Arp2/3 complex (Konigs *et al*, 2014; Qi & Yu, 2025; Rotty *et al*, 2017; Wells *et al*, 2004), all of which are essential for efficient IL-4-induced fusion (Balabiyev *et al*, 2020; Faust *et al*., 2019; Jay *et al*., 2007). Transcriptomic analysis revealed downregulation of genes encoding plasma membrane and cortical components, along with altered actin turnover and misregulated activity of Cofilin and WASP. In IL-4-stimulated *control* macrophages, both WASP and Filamin A were enriched in the nucleus, consistent with their reported non-canonical functions in transcriptional regulation and chromatin remodeling (Wu *et al*, 2006; Yuan *et al*, 2022). In *Atf3*-deficient cells, WASP remained cytoplasmic, and Filamin A accumulated in perinuclear foci. Since WASP nuclear entry depends on Src family kinase-mediated phosphorylation downstream of Cdc42 (Looi *et al*, 2014), changes in WASP localization and levels despite reduced F-actin network suggest impaired Rho GTPase activation. Given that both WASP and Filamin A are essential for podosome formation and function (Faust *et al*., 2019; Guiet *et al*, 2012; Song *et al*, 2014), their misregulation likely contributes to defective spreading and fusion observed in *Atf3* deficient cells.

We link these cytoskeletal defects to dysregulated lipid metabolism. Loss of *Atf3* leads to elevated *Ch25h* expression and accumulation of its oxysterol product 25-hydroxycholesterol (25-HC). 25-HC suppresses the mevalonate pathway, reducing cholesterol levels, altering membrane composition, and decreasing isoprenoid intermediates. This, in turn, impairs prenylation of Rho GTPases, disrupting actin remodeling and weakening the coordination between the plasma membrane and the actin cytoskeleton that is essential for spatially and temporally controlled membrane dynamics during cell-cell fusion. The similarity between phenotypes induced by *Atf3* deficiency, 25-HC treatment, and pharmacological inhibition of prenylation, together with partial rescue of actin dynamics upon *Ch25h* deletion, highlights the importance of integrated lipid-cytoskeletal regulation in IL-4-driven fusion. While *Ch25h* is best known as an interferon-inducible regulator of cellular cholesterol content and inflammasome activation (Adams *et al*, 2004; Dang *et al*, 2017; Park & Scott, 2010; Reboldi *et al*, 2014), 25-HC also inhibits virus-cell fusion by depleting an accessible pool of plasma membrane cholesterol (Heisler *et al*, 2023; Liu *et al*, 2013; Wang *et al*, 2020; Zang *et al*, 2020). Our findings extend this function of 25-HC to macrophage fusion. Notably, *Ch25h* expression is transiently induced in IL-4-stimulated macrophages (Czimmerer *et al*, 2018, our data - Supplementary Dataset 1) suggesting a physiological role for 25-HC in fine-tuning immune-metabolic responses during early alternative activation. However, sustained 25-HC elevation, as in Atf3 deficient macrophages, shifts this balance toward fusion suppression, a mechanism paralleled by statins, which act as competitive inhibitors of Hmgcr, thereby lowering cholesterol synthesis, but also intermediate products of the mevalonate pathway, including isoprenoids. Statins, like 25-HC, affect Rho GTPases-mediated signaling and actin network with impact on macrophage adhesion and motility (Ako *et al*, 2022; Akula *et al*., 2019; Healy *et al*, 2020; Kuipers *et al*, 2006).

### Ch25h-independent control of nuclear lamina integrity

Restoring cholesterol levels and actin dynamics in *Atf3*^*KO*^*Ch25h*^*KO*^ macrophages did not rescue fusion, revealing a Ch25h-independent role for Atf3. *Atf3*-deficient macrophages showed markedly reduced lamin A/C expression, nuclear deformation, increased fragility, and elevated DNA damage. Given the mechanical coupling between cytoskeleton and nucleoskeleton, lamin A/C depletion may impair force transmission during later stages of the fusion process, affecting membrane apposition or fusion pore formation. Impaired multinucleation of *Lmna*^*G609G*^ BMDMs provides independent support for a requirement of intact cyto-nucleosleketal mechanotransduction. Although the *Lmna*^*G609G*^ model involves progerin accumulation and therefore does not phenocopy simple lamin A/C depletion, the observed fusion defect is consistent with the notion that an intact nuclear lamina contributes to efficient macrophage multinucleation.

The persistent reduction of *LMNA* mRNA and lamin A/C protein irrespective of Ch25h status is compatible with direct transcriptional regulation by Atf3, in line with its control of the *Drosophila* ortholog (Donohoe *et al*., 2018). Nevertheless, we cannot exclude the contribution of altered lamin post-translational processing, such as impaired farnesylation, in *Atf3*-deficient cells. The DNA damage signature observed in *Atf3*^*KO*^ BMDMs further aligns with the genomic instability observed in *Atf3*-deficient mouse embryonic fibroblasts (Wang *et al*, 2018), supporting a broader role for Atf3 in maintaining genome integrity, although the underlying mechanisms are likely context dependent.

### Implications and future directions

Atf3 emerges as a central regulator of macrophage fusion competence in IL-4 dominated environments, coupling lipid-cytoskeletal regulation via Ch25h repression with lamin A/C-dependent nuclear stability. This dual control ensures both membrane plasticity and nuclear resilience required for giant cell formation. Disruption of either pathway impairs fusion while combined loss, as observed in *Atf3* deficiency, blocks it entirely. Our findings have direct relevance for conditions involving macrophage multinucleation, including foreign body reactions and atherosclerosis, where Atf3 has been implicated (De Nardo *et al*., 2014; Gold *et al*., 2012; and this study), as well as osteoclastogenesis, or granulomatous inflammation, in which coordination of mechanotransduction and lipid metabolism may occur independently of Atf3. The inhibitory effect of sustained 25-HC parallels that of statins, widely used cholesterol lowering drugs, raising the possibility that pharmacological or endogenous suppression of the mevalonate pathway converges on shared mechanism limiting giant cell formation. These suggests that statin therapy may alter FBGC responses or osteoclastogenesis, with implications for biomaterial integration and bone homeostasis. Similarly, defects in lamin A/C, as seen in laminopathies or during natural aging (McClintock *et al*, 2007; Scaffidi & Misteli, 2006) may compromise nuclear mechanics in other fusion-dependent contexts, including muscle regeneration to placental development. Modulating Atf3 or its downstream effectors may therefore represent a strategy to fine-tune multinucleated giant cell formation in therapeutic settings.

## METHODS

### Generation, breeding and genotyping of mutant and transgenic mouse strains

Animals were housed in specific-pathogen-free (SPF) conditions in the *in vivo Research Facility* (ivRF) of the *CECAD* Research Center and monitored by routine testing of sentinel mice. Mice had *ad libitum* access to drinking water and normal chow diet (R/M-H-V1554, ssniff Spezialdiäten GmbH) containing 55.1% carbohydrates, 19.3% proteins, and 3.3% fat (9% calories from fat). Mouse breeding and handling were conducted in accordance with the ivRF guidelines, and under approval of the Animals Ethics Committee of North Rhine-Westphalia, Germany (AZ 84-02.04.2016.A318). The *Atf3*^*KO*^ (*Atf3*^*-/-*^) and *Ch25h*^*KO*^ (*Ch25h*^*-/-*^) full body knock-out mouse strains were generated using the CRISPR/Cas9 technique in the transgenic core unit of the ivRF by implanting the one-cell stage fertilized oocytes injected with Cas9-sgRNA mixture to pseudo-pregnant C57BL/6N mice. To generate *Atf3*^*KO*^ strain, two guide RNAs targeting exon 2 and exon 4 of the mouse Atf3 locus (Chr1:191170296-191183333 bp, - strand), respectively, were used to generate a deletion of 5933 bp, including the bZIP domain. Full body loss of function *Ch25h*^*KO*^ mice were generated by targeting the single exon of the *Ch25h* gene (Chr19: 34451183-34452548 bp, - strand) with two guide RNAs causing a 49 bp deletion, which led to a frame shift in the coding sequence and a premature stop codon. Of note, the 49 bp deletion also affects the first intron of the long noncoding RNA *Gm26902* coded on the opposite strand. The intronic deletion should not affect *Gm26902* expression nor processing. *Atf3*^*KO*^ and *Ch25h*^*KO*^ lines were backcrossed to the C57BL/6 genetic background for several generations before carrying out experiments. Heterozygous *Atf3*^*+/-*^ and *Ch25h*^*+/-*^ animals were further intercrossed to generate *Atf3*^*KO*^*Ch25h*^*KO*^ (*Atf3*^*-/-*^ and *Ch25h*^*-/-*^) mice. *Lifeact-EGP, Rosa26, mTmG-TdTOMATO* and *H2B-EGFP* mice were kindly provided by Prof. Dr. Carien Niessen (CECAD, University of Cologne), Dr. Claudia Dafinger (CECAD, University of Cologne), and Prof. Dr. Sandra Iden (Faculty of Medicine & ZHMB, Saarland University), respectively. The three lines were crossed to *Atf3*^*KO*^ and *Ch25h*^*KO*^ mutant lines. A *Lmna*^*G609G*^ knock-in mouse strain (Osorio *et al*., 2011) was housed at University of Oviedo animal facility and monitored by routine testing of sentinel mice. Mice were caged separately by sex in cages with solid floors, sawdust and nests. Mice had *ad libitum* access to drinking water and normal chow diet (SAFE® A40, SAFE) containing 60.4% carbohydrates, 15.2% proteins, and 3.2% fat. Three times a week, mice were given moistened ground pellets of food to facilitate the feeding of progeroid mice. All components of the cages, including food, had been autoclaved previously. Mouse breeding and handling were conducted in accordance with the guidelines of the Committee for Animal Experimentation of the Universidad de Oviedo, and under the approval of the Regional Ethics Committee for Animal Experimentation (PROAE 02/2023) For all experiments, homozygous *Lmna*^*G609G*^ 8-12-week-old mice were used, and compared to age-matched, gender-matched control mice when possible. The sequence of targeting gRNAs and genotyping primers are listed in Supplementary Table 1.

### Isolation, handling and treatment of bone marrow-derived macrophages (BMDMs)

Bone marrow cells were isolated as previously described (Trouplin *et al*, 2013) and differentiated for 6 days in DMEM 1x + GlutaMax-I medium (Gibco #31966-021) supplemented with 10% fetal Bovine Serum (FBS) Superior (Biochrom #S0615), 1% penicillin/streptomycin (10,000 U/mL) (Thermo Scientific #15140-122), 1% HEPES (1 M) (Thermo Scientific #15630106) and 50 ng/mL recombinant mouse Macrophage Colony-Stimulating Factor (rm M-CSF, ImmunoTools #12343117). On day 6, differentiated BMDMs were seeded at a density of 0.75 × 10^5^ cell/well in 8-well glass chamber slides (Idibi #80827) or Permanox slides (Thermo Scientific #177445) for confocal microscopy; 1 × 10^5^ BMDMs in 35 mm dishes for spinning disk live imaging; 1 × 10^6^ cells/well in 6-well plates for RNA and protein analyses. After cells adhered, BMDMs were treated with medium containing only rm M-CSF (25 ng/mL) (unstimulated, unst.) or in combination with recombinant murine IL-4 (Peprotech #214-14) at a concentration of 25 ng/mL for 3 days, unless indicated otherwise. For fusion assays, 3.75 × 10^4^ cell/well of BMDMs expressing *Rosa26, mTmG-TdTOMATO* were mixed with 3.75 × 10^4^ cell/well of cells expressing H2B-EGFP in 8-well glass chamber slides. Osteoclasts were differentiated from BMDMs in medium containing 25 ng/mL rm M-CSF and 50 ng/ml RANKL (Peprotech #315-11) for 9 days. The medium was exchanged every 3 days. TRAP staining of osteoclasts was performed using Acid Phosphatase, Leukocyte (TRAP) Kit (Sigma-Aldrich #387A). For the TLR2-stimulated multinucleation assays, FSL-1 (Invivogen #tlrl-fsl) and Pam3CSK4 (Invivogen #tlrl-pms) were used at a concentration of 100 ng/mL, together with rm M-CSF (25 ng/mL). For the *Mycobacterium tuberculosis* infection assay, BMDMs were seeded in medium without antibiotics and supplemented with rm M-CSF (25 ng/mL). Infection was performed at a multiplicity of infection equal 1 for the Erdman strain (ATCC 35801). Treatment of BMDMs with 25-HC (2.5 ng/mL) (Sigma #H1015), Lonafarnib (5 µM and 10 µM) (Sigma Aldrich #SML1457) and GGTI-2133 (5 µM and 10 µM) (Sigma Aldrich #G5294) was carried out in rm M-CSF (25 ng/mL) supplemented medium for 24 hours before treatment with IL-4 (25 ng/mL) for 3 days.

### Isolation, handling and treatment of peritoneal macrophages (PMs)

Peritoneal Macrophages (PMs) were isolated from the peritoneal cavity of 8-10-week-old mice according to a previously described protocol (Herb *et al*, 2019) using paramagnetic CD11b MicroBeads (MiltenyiBiotec #130-049-601). After isolation, PMs were seeded at 1.5 × 10^5^ cells/well in 8-well glass chamber slides or Permanox slides and stimulated with IL-4 (25 ng/mL) for 6 days.

### Immunostaining

Cells were washed three times in PBS and fixed in 4% paraformaldehyde (PFA) in PBS for 25 min at RT. Following fixation, cells were washed three times in PBS, permeabilized in PBS containing 0.5% Triton X-100 (PBS-T) for 10 min and incubated in a blocking buffer (3% BSA in PBS) for 20 min at RT. Primary antibodies were diluted in DAKO-background reducing solution (Agilent #S3022) and incubated with cells overnight at 4°C. The following primary antibodies were used: anti-GFP (goat, 1:500; Abcam #ab6673, RRID:AB_305643), anti-CD11b (rat, 1:200; DSHB #M1/70.15.11.5.2, RRID:AB_2234066), anti-Phospho-Myosin Light Chain 2 (Ser19) (rabbit, 1:200; Cell Signaling Technology #3671, RRID:AB_330248), anti-WASP (mouse; 1:200; Santa Cruz #sc-13139, RRID:AB_628445), anti-Filamin A (rabbit, 1:200; Bethyl-Biomol #A301-135A), anti-lamin A/C (mouse, 1:250, Cell Signaling Technology #4777, RRID:AB_10545756). Cells were washed three times in blocking buffer and incubated for 1 hour with corresponding secondary antibodies (1:500): Cy3 AP Anti-Mouse IgG (Jackson ImmunoResearch #715-165-151; RRID:AB_2315777), Cy5 AP Anti-Mouse IgG Cy3 AP (Jackson ImmunoResearch #715-175-151; RRID:AB_2340820), Cy3 AP Anti-Rabbit IgG (Jackson ImmunoResearch Labs #711-165-152, RRID:AB_2307443), Cy5 AP Anti-Rabbit IgG (Jackson ImmunoResearch #711-175-152; RRID:AB_2340607), Cy2 AP Anti-Goat IgG (Jackson ImmunoResearch Labs #705-225-147, RRID:AB_2307341), Cy3 AP Anti-Rat IgG (Jackson ImmunoResearch Labs #712-165-150, RRID:AB_2340666), Cy5 AP Anti-Goat IgG (Jackson ImmunoResearch Labs #712-175-153, RRID:AB_2340672), 4’,6-Diamidino-2-phenylindol Dihydrochloride (DAPI) (1 µg/mL, Carl Roth GmbH #6335) and Alexa Fluor 488 or 546 Phalloidin (1:1000, Thermo Scientific #A12379, #A22283). After staining, cells were washed three times in PBS then briefly in dH_2_O before mounted in Mowiol-DABCO solution. See Supplementary Table 1 for details on antibodies.

### Image acquisition and processing

Bright-field images were acquired using an Olympus CKX41 microscope equipped with a DP72 camera and CellSens 1.1 software. Bright-field time-lapse live images were acquired using a JuLI Smart fluorescent image analyzer with a 10x objective or an EVOS FL Auto 2 microscope (Thermo Scientific) equipped with a Plan S-Apo 20x (NA 0.75) objective. Time-lapse live imaging of fluorescently labeled cells was performed using either an UltraView VoX spinning disk confocal microscope (Perkin Elmer) equipped with Plan-Fluor 20x (NA 0.75 Oil), Plan-Fluor 40x (NA 1.3 Oil), and Plan-Apo Tirf 60x (NA 1.49 Oil) objectives, or a Zeiss LSM980 Airyscan 2 equipped with a Plan-Apo 63x (NA 1.4 Oil DIC) objective. Confocal images of fixed samples were acquired at room temperature using an Olympus FV1000 confocal microscope equipped with 40x UPlan FL (NA 1.30) and 60x UPlanApo (NA 1.35) objectives and a Leica SP8 confocal system equipped with PL Apo 63x (NA 1.20 water) and HC PL APO 93X (NA 1.30 GLYC) objectives. Acquired z-stacks were deconvoluted using Huygens Software (Scientific Volume Imaging, SVI). Transversal sections, z-stacks and time-lapse images were generated and processed using ImageJ (RRID:SCR_002285). Panel assembly and brightness/contrast adjustments were done using Adobe Photoshop CC (Adobe Systems Inc., RRID:SCR_014199). To calculate the Fusion Index (FI), the total number of nuclei within MGCs was divided by the total number of nuclei per image. The FI is expressed in %. Cell “roundness” (roundness = 4A/(π*Major Axis^2)) and nuclear “circularity” (circularity = 4π(area/perimeter^2)) were calculated in ImageJ. Images used for quantification of fluorescence intensity were acquired using the same settings. WASP fluorescence intensity was calculated using the function “Integrated density” in ImageJ on single cells.

### Electron microscopy

Cells were seeded on Permanox chamber slides. After two PBS washes, cells were fixed with 2.5% EM-grade glutaraldehyde for 2 hours, washed in phosphate buffer (pH 7.2), and post-fixed with 2% osmium tetroxide in phosphate buffer for 30 - 60 minutes at RT in the dark. Cells were then washed in phosphate buffer, followed by dH_2_O washes. Gradual dehydration was performed on ice using an ethanol series (30%, 50%, 70%, 90%, 96%, 5 minutes each), followed by two 10-minute washes in 100% ethanol and two in 100% acetone at RT. The slides were incubated overnight at 4°C in an acetone (1:1) mixture. Acetone was evaporated under a hood for 1.5 hours, then replaced with fresh araldite. After 4 hours, araldite was partially removed, and fresh araldite was added to fill half the well, followed by polymerization at 65°C. After 24 hours, polymerized cell blocks were separated from the slides. Semi-thin (1 µm) sections were stained with toluidine blue, and thin (70–80 nm) sections were stained with 2% uranyl acetate and lead citrate. Thin sections were imaged with a Zeiss EM 109 transmission electron microscope at 80 kV.

### Microchannels

PDMS microchannels inserted in Petri dishes were purchased from 4D Cell (#MC011, #MC019). The dishes were coated with fibronectin (25 µg/mL) according to manufacturer’s instructions and incubated with pre-warmed medium containing rm M-CSF (25 ng/mL) for 15 minutes at 37°C, prior to cell seeding. Differentiated BMDMs were seeded in high confluency (4 × 10^4^ cells/ access port) and allowed to adhere for at least 1 hour (37°C, 5% CO_2_). Four hours prior to live imaging, medium with IL-4 (25 ng/mL) was added. For immunostaining, microchannels were washed in PBS and BMDMs were fixed by adding 4% PFA solution to each access port for 30 minutes. PDMS was carefully removed from the dishes, and the immunostaining protocol was applied on the surface in contact with the cells. For imaging of the fixed and stained BMDMs, PDMS microchannels were placed on 35 mm imaging dishes (Ibidi #81156).

### RNA isolation and quantitative reverse transcription PCR (RT-qPCR)

BMDMs were washed twice with PBS, collected in 800 µL of TRI reagent (Sigma-Aldrich, #T9424) containing β-mercaptoethanol (10 µL/mL), and snap-frozen in liquid nitrogen. After adding 200 µL of chloroform, samples were vigorously mixed and centrifuged for 15 minutes at 12,000 × g (4°C), and the upper phase was collected. An equal volume of isopropanol was added; samples were vortexed for 5 seconds and transferred to a ReliaPrep™ Minicolumn. The next steps of RNA isolation followed the protocol of the ReliaPrep™ RNA Cell Miniprep System Kit (Promega #z6011), including DNAse I treatment (Promega #Z358A) to minimize genomic DNA contamination. cDNA was synthesized from 1 to 2 µg of RNA using oligo (dT) primers and the reverse transcriptase. The RT-qPCR was performed in technical triplicates with at least three independent biological replicates per genotype with GoTaq polymerase mix (Promega #A6002) in a CFX96 real time PCR system (Biorad), based on SYBR Green dye or probe detection. All primers were designed to anneal at 62°C. See Supplementary Table 1 for primer sequences.

Expression values were normalized to the levels of the housekeeping gene *Gapdh*, and fold-changes in gene expression were calculated using the ΔΔC_T_ method (Livak & Schmittgen, 2001). For *Atf3* levels, GoTaq probe qPCR master mix was used with pre-designed probe, *Atf3*-FAM (IDT). Data were normalized to the levels of the housekeeping *Gapdh*-HEX probe present in the same reaction, using the above-mentioned method. Statistical significance of gene expression changes was determined using Student’s t-test assuming unequal variance (Mann-Whitney test) or one-way ANOVA with Holm-Šídák’s or Tukey’s correction for multiple comparisons as specified in the corresponding Figure legend.

### Differential gene expression and analysis

Total RNA was isolated as described above with four biological replicates per experimental group. For RNA sequencing, 2 μg of RNA were used for library preparation (Illumina TruSeq Stranded total RNA Ribo-Zero). Pair-end sequencing (100 bp) was performed using the Illumina HiSeq 2000 platform. Pseudoalignment of reads to the total *Mus musculus* cDNA sequences (GRCm39_107) was done by kallisto v0.46.1 (Bray *et al*, 2016) (RRID:SCR_016582). Differential gene expression was determined using *DESeq2* v1.34.0 (Love *et al*, 2014) (RRID:SCR_015687). Heat maps comparing expression levels of selected genes between samples were constructed from DESeq2 normalized counts and log_2_FC values using R-studio. Gene Ontology term enrichment analysis was performed using *ShinyGO* v0.82 (Ge *et al*, 2020) using a custom background gene list containing all genes detected by the RNA-seq analysis. Fisher’s exact test with false discovery rate (FDR) correction was used.

### Preparation of samples for western blotting

PBS-washed cells were scraped and immediately lysed in RIPA lysis buffer (50 mM HCl pH 8, 150 mM NaCl, 1% NP-40, 0.5% sodium deoxycholate, 0.1% SDS, protease and phosphatase inhibitors). All samples were cleared of debris and DNA by centrifugation (18,000 x g at 4°C for 10 min). Protein concentration was determined by Bradford assay (Roth GmbH K015.1) according to manufacturer’s instructions. Samples were denatured by boiling in Laemmli buffer containing 10% β-mercaptoethanol for 5 minutes at 95°C. After resolving on 10% or 12% SDS-PAGE, proteins were transferred onto a nitrocellulose membrane, followed by one hour blocking at RT in Tris Buffered Saline buffer containing 0.1% Tween and 3% BSA. Candidate proteins were detected by immunoblotting with primary antibodies overnight at 4°C. The following primary antibodies were used: anti-ATP5a [15H4C4] (mouse, 1:1000; Abcam #ab14748, RRID:AB_301447), anti-ATF3 (rabbit, 1:500, Novus #NBP1-85816, RRID:AB_11014863), anti-actin (rabbit, 1:1000, Sigma-Aldrich #A2066, RRID:AB_476693), anti-STAT6 (rabbit, 1:1000, Cell Signaling Technology #5397, RRID:AB_11220421), anti-phospho-STAT6 (Tyr641) (D8S9Y) (rabbit, 1:1000, Cell Signaling Technology #56554, RRID:AB_2799514), anti-STAT1 (rabbit, 1:1000; Cell Signaling Technology #9172, RRID:AB_2198300), anti-phospho-STAT1 (Tyr701) (58D6) (rabbit, 1:1000, Cell Signaling Technology #9167, RRID:AB_561284), anti-WASP (mouse; 1:500; Santa Cruz #sc-13139, RRID:AB_628445), anti-lamin A/C (mouse, 1:1000, Cell Signaling Technology #4777, RRID:AB_10545756), anti-Phospho-Cofilin (Ser3) (77G2) (rabbit, 1:1000; Cell Signaling Technology #3313, RRID:AB_2080597), anti-Cofilin (D3F9) (rabbit, 1:1000; Cell Signaling Technology #5175, RRID:AB_10622000), anti-HMGCR (mouse, 1:500, Atlas Antibodies #AMAb90619, RRID:AB_2665607), anti-HMGCS1 (mouse, 1:500; Novus #NBP2-36554, RRID:AB_3295882). Following five washes (TBS, 0.1% Tween), membranes were incubated with corresponding secondary HRP-conjugated antibodies (1:5000) at RT for 2 hours. The following secondary antibodies were used: HRP AP Anti-Mouse IgG (Jackson ImmunoResearch Labs #715-035-150, RRID:AB_2340770), HRP AP Anti-Rabbit IgG (Jackson ImmunoResearch Labs #711-035-152, RRID:AB_10015282). After several washes in TBS, 0.1% Tween, chemiluminescent signal was developed with ECL Western blotting substrate (Promega W1015) and captured using ImageQuant LAS4000 reader (GE Healthcare, RRID: SCR_014246). See Supplementary Table 1 for details on antibodies.

### Quantification of protein signals in immunoblot analysis

Signal intensities were quantified using the Gel Analyzer tool in ImageJ (https://fiji.sc/) (RRID: SCR_003070). The intensity of the protein of interest was normalized to the intensity of the respective loading control band for each biological replicate. Phosphorylated protein levels are presented as relative expression in relation to the sum of normalized total and phosphorylated band intensities. Statistical significance for changes in protein levels was determined using an unpaired Student’s t-test or one-way ANOVA followed by Tukey’s multiple comparisons test.

### G- and F-actin fractionation and quantification

The F- to G-actin ratio was determined as previously described (Le *et al*, 2016). Cells were harvested in cytoskeleton-stabilizing lysis buffer (50 mM PIPES pH 6.9, 50 mM NaCl, 5 mM MgCl_2_, 5 mM EGTA, 0.2 mM dithiothreitol, 0.1% NP-40, 0.1% Tween-20, 5% glycerol, 1 mM ATP, and protease inhibitors). After ultracentrifugation at 55,000 rpm (TLA-55 rotor) for 1 hour at 4°C, the supernatant was collected as the G-actin fraction. The pellet was solubilized in actin-depolymerizing buffer (50 mM PIPES pH 6.9, 5 mM MgCl_2_, 10 mM CaCl_2_, 5 µM cytochalasin D), sonicated for 5 minutes, and used as the F-actin fraction. Protein concentration was determined by the Bradford assay (Roth GmbH, K015.1) according to the manufacturer’s instructions. Concentrations were normalized between samples from the same actin fraction. Lysates were then processed for western blotting as described above. Equal loading was verified by Ponceau staining of the membrane. F/G actin levels are represented as a ratio of F-actin under G-actin levels. Statistical significance was determined using ordinary one-way ANOVA with a post hoc Tukey’s multiple comparison test.

### Lipid measurements in BMDMs

BMDMs (15 × 10^6^ cells) stimulated with IL-4 (25 ng/mL) for 24 hours were washed with ice-cold PBS and collected for measurement of selected lipid species. Total cellular cholesterol was determined using Liquid Chromatography/Mass Spectrometry, while Glycerophospholipids (GPLs) and Cholesterol Esters (CE) were measured by Shotgun lipidomics at the CECAD Lipidomics/Metabolomics Core Facility. Levels were normalized to protein content. Statistical significance was determined using ordinary one-way ANOVA with Holm-Šídák’s multiple comparison test.

### Quantification and statistical analysis

All statistical analyses were performed in Graphpad Prism version 10, as defined throughout the text, materials and methods and Figure legends.

## FUNDING

This work was funded under Germany’s Excellence Strategy – CECAD, EXC 2030 – 390661388 from the Deutsche Forschungsgemeinschaft (DFG, German Research Foundation). DFG grants UH 243/8-1 - Projektnummer: 573567388 to M.U. and SPP2493 ID79/7-1 - Projektnummer 564825595 to S.I. Funding for instrumentation: Zeiss LSM980 Airyscan 2 (DFG-INST 216/1068-1FUGG); Leica SP8 confocal system (DFG-INST 2016/742-1 FUGB). The *Lmna*^*G609G*^ mice housing was funded by the Ministerio de Ciencia, Innovación y Universidades (Spain) (PID2023-148089OB-I00).

## ACKNOWLEDGEMENTS

We thank Manolis Pasparakis, Carien Niessen, Claudia Dafinger, and their labs for advice, sharing antibodies, mouse strains and reagents. We thank Jan Rybniker and Jessica Graeb for advice and help with conducting *Mycobacterium* infection experiments, Michael Schramm for help with isolation of peritoneal macrophages, Ferdinand Grawe for electron microscopy, and Susanne Brodesser from the CECAD Lipidomics/Metabolomic Facility for lipid measurements. We thank Maria Asif for help with BMDM isolation and culturing. We are grateful to Katja Brückner, Katrin Kierdorf, Hisham Bazzi for technical advice, project discussions, critical reading and comments on the manuscript, Bianca Collins, Agnieszka Sokol, Nils Teuscher, Maria Serradas, Je Cuan Choo, Susane Kolek, and Jose Luis Rodriguez Garcia for excellent technical assistance. We thank the CECAD *in vivo* research facility for generation and care of the *Atf3*^*KO*^ and *Ch25h*^*KO*^ mice and José M. P. Freije for sharing the tissues from *Lmna*^*G609G*^ mouse line. We are grateful to the CECAD Imaging facility for helping with setting up various imaging techniques.

## AUTHORS CONTRIBUTIONS

A.C. and M.U. originated the initial idea. A.C., M.U. and N.M. jointly formulated the concept. A.C., C.J., E.K.W., M.D.G, M.U., and N.M. carried out the investigation. A.C., D.S., E.K.W., G.C., M.U., N.M. and S.I. performed formal analyses. A.G.T provided tissues from *Lmna*^*G609G*^ mice and the respective controls. A.C., N.M. wrote the initial draft. M.U. wrote the manuscript and generated the visualizations therein. E.K.W., G.C., S.I. and M.U. edited the manuscript. M.U. provided supervision, project administration, and funding.

## CONFLICT OF INTEREST DISCLOSURE

The authors declare no competing interests.

## DATA AVAILABILITY

Original data and any additional information required to reanalyze the data reported in this paper will be shared by MU upon request. The datasets produced in this study are available in the following databases:

RNA-Seq data: Gene Expression Omnibus (RRID:SCR 007303) (Edgar et al., 2002) GSE240477 (https://www.ncbi.nlm.nih.gov/geo/query/acc.cgi?acc=GSE240477).

---

**Supplementary Movie 1. IL-4-induced multinucleation of control BMDMs involves both acytokinetic division and cell-cell fusion**.

Bright field time-lapse live imaging of *ctrl* BMDMs stimulated with IL-4 for three days shows incomplete cytokinesis (rectangle) and cell-cell fusion (asterisks). See Figure 2B for video stills.

**Supplementary Movie 2. Actin cytoskeleton dynamics in IL-4-stimulated control BMDMs**.

Live-cell time-lapse confocal microscopy of IL-4-stimulated *ctrl* BMDMs expressing Lifeact-EGFP shows dynamic rearrangement of the actin cytoskeleton, including podosome (asterisks), lamellipodia (arrowhead), and filopodia (arrow) formation, as well as membrane ruffling. Scale bar: 10 µm. See Figure 4K for video stills.

**Supplementary Movie 3. Actin cytoskeleton dynamics in IL-4-stimulated Atf3**^***KO***^ **BMDMs**.

Live-cell time-lapse confocal microscopy of IL-4-stimulated *Atf3*^*KO*^ BMDMs expressing Lifeact-EGFP reveals an elongated, stellate morphology, with reduced actin cytoskeleton dynamics, limited membrane ruffling and diminished podosome formation. Scale bar: 10 µm. See Figure 4L for video stills.

**Supplementary Movie 4. Nucleus of IL-4-stimulated control BMDMs traversing through constriction**. Live-cell time-lapse confocal microscopy of IL-4-stimulated *control* cells expressing H2B-GFP as they pass through 1.5-2 µm constrictions. Scale bar: 10 µm. See Figure 7R for video stills.

**Supplementary Movie 5. Nucleus of IL-4-stimulated Atf3**^*KO*^ **BMDMs traversing through constriction**. Live-cell time-lapse confocal microscopy of IL-4-stimulated *control* cells expressing H2B-GFP as they pass through 1.5-2 µm constrictions with nucleo-cytoplasmic GFP leakage being observed over time. Scale bar: 10 µm. See Figure 7S for video stills.

**Supplementary Dataset 1**. RNA-seq results (differential gene expression of unstimulated and 5-hour IL-4-stimulated control and Atf3-deficient bone marrow derived macrophages).

**Supplementary Dataset 2**. RNA-seq results (GO term enrichment analysis of unstimulated and 5-hour IL-4-stimulated control and Atf3-deficient bone marrow derived macrophages).

## Notes

https://www.ncbi.nlm.nih.gov/geo/query/acc.cgi?acc=GSE240477

### Competing Interest Statement

The authors have declared no competing interest.

