## Supplementary Table S1 for "Atf3 Integrates Lipid and Cytoskeletal Remodeling to Drive Macrophage Fusion"

**Supplementary Table S1. Experimental reagents, software and equipment**

**Targeting single guide oligonucleotides/RNAs**

| Oligo | Sequence (5′‐ 3′) |
| --- | --- |
| Atf3 Exon 2 IVT sgRNA | GAAA**TTAATACGACTCACTATAGG**AGCGCAGAGGACATCCGAGTTTTAGAGCTAGAAATAGC |
| Atf3 Exon 4 IVT sgRNA | GAAA**TTAATACGACTCACTATAGG**CGGACACCGGAAGACGAGGTTTTAGAGCTAGAAATAGC |
| sgRNAR | AAAAGCACCGACTCGGTGCCACTTTTTCAAGTTGATAACGGACTAGCCTTATTTTAACTTGCTATTTCTAGCTCTAAAAC |
| Ch25h sgRNA 1 | /AlTR1/rCrArUrGrGrGrCrUrGrCrUrArCrArArCrGrGrUrUrGrUrUrUrUrArGrArGrCrUrArUrGrCrU/AlTR2/ |
| Ch25h sgRNA 2 | /AlTR1/rGrGrUrUrCrGrGrArGrCrUrCrCrArArGrArCrCrUrGrUrUrUrUrArGrArGrCrUrArUrGrCrU/AlTR2/ |

**Screening and genotyping primers**

| Primer | Sequence (5′‐ 3′) |
| --- | --- |
| #2761 Atf3 Intron 2 Rev | CACACTCTATCTGCCCATTCTC |
| #2178 Atf3 Exon 2 Surv For | GGGCATGGGCATAAAGTAATAAG |
| #2181 Atf3 Exon 4 Surv Rev | CAGAGTGGGTCAATGCAGTAG |
| #2431 Ch25h del For | GCCACAGTCTTAAGAAAAGTCGGC |
| #2432 Ch25h del Rev | ACGCCTCCCTTGTCCTTATGG |
| #3169 Rosa26 mTmG-TdTOMATO-For | CGAGGCGGATCACAAGCAATA |
| #3170 Rosa26 mTmG-TdTOMATO-Rev | CGAGGCGGATCACAAGCAATA |
| #3171 Rosa26 mTmG-TdTOMATO-Rev | TCAATGGGCGGGGGTCGTT |
| #1937 H2BGFP genotyp JAX | TCCTTGAAGAAGATGGTGCG |
| #1938 H2BGFP genotyp JAX | CTAGGCCACAGAATTGAAAGATCT |
| #1939 H2BGFP genotyp JAX | GTAGGTGGAAATTCTAGCATCATCC |

**RT-qPCR primers**

| Gene | Primer sequence (5′‐ 3′) forward | Primer sequence (5′‐ 3′) reverse |
| --- | --- | --- |
| *Atf3* | GGAGATGTCAGTCACCAAGTCTGAG | CAGTTTCTCTGACTCTTTCTGCAGG |
| *Il6* | AACAGAGACGCATGATGTGCTCCA | AATCGGTTACGTTCTGCAAGTCGC |
| *Ch25h* | CCAGCTCCTAAGTCACGTC | CACGTCGAAGAAGGTCAG |
| *Isg15* | TGGGACCTAAAGGTGAAGATGCTG | TGCTTGATCACTGTGCACTGGG |
| *Hmgsc1* | CCTATGATTGCATTGGGCGGC | CATCCCAAGAGCTGGACTCAA |
| *Hmgcr* | TCCCACCCCTGGGAAGTTATT | AGGATGATGATGTCGCTGCTC |
| *Lmna* | AAGACCCTTGATTCTGTGGC | TCCAATGTGCGCTTCTCAC |
| *Gapdh* | ACAGTCCATGCCATCACTGCC | GCCTGCTTCACCACCTTCTTG |
| GoTaq probes (Predesigned qPCR Assays, IDT) | | **Assay ID** |
| *Gapdh* | | Mm.PT.39a.1 |
| *Atf3* | | Mm.PT.58.41654515 |

**Immunostaining**

| Antibody/Dye | Source | RRID | Catalog Number |
| --- | --- | --- | --- |
| αTubulin | Developmental Studies Hybridoma Bank | RRID:AB_579793 | #AA-4.3 |
| CD11b | Developmental Studies Hybridoma Bank | RRID:AB_2234066 | #M1/70.15.11.5.2 |
| Filamin A | Bethyl-Biomol |  | #A301-135A |
| lamin A/C | Cell Signaling Technology | RRID:AB_10545756 | #4777 |
| WASP | Santa Cruz | RRID:AB_628445 | #sc-13139 |
| Phospho-Myosin Light Chain 2 (Ser19) | Cell Signaling Technology | RRID:AB_330248 | # 3671 |
| GFP | Abcam | RRID:AB_305643 | #ab6673 |
| Phospho-γH2A.X (Ser139) | Cell Signaling Technology | RRID:AB_2118009 | #9718 |
| Alexa Fluor 488 Phalloidin | Thermo Scientific |  | #A12379 |
| Alexa Fluor 546 Phalloidin | Thermo Scientific |  | #A22283 |
| 4’,6-Diamidino-2-phenylindol Dihydrochlorid (DAPI) | Carl Roth GmbH |  | #6335 |
| Cy3 AP Anti-Mouse IgG (H+L) | Jackson Immunoresearch | RRID:AB_2315777 | #715-165-151 |
| Cy5 AP Anti-Mouse IgG (H+L) | Jackson Immunoresearch | RRID:AB_2340820 | #715-175-151 |
| Cy3 AP Anti-Rabbit IgG (H+L) | Jackson Immunoresearch | RRID:AB_2307443 | #711-165-152 |
| Cy5 AP Anti-Rabbit IgG (H+L) | Jackson Immunoresearch | RRID:AB_2340607 | #711-175-152 |
| Cy3 AP Anti-Rat IgG (H+L) | Jackson Immunoresearch | RRID:AB_2340666 | #712-165-150 |
| Cy2 AP Anti-Goat IgG (H+L) | Jackson Immunoresearch | RRID:AB_2307341 | #705-225-147 |
| Cy5 AP Anti-Goat IgG (H+L) | Jackson Immunoresearch | RRID:AB_2340672 | 712-175-153 |

**Immunoblotting**

| STAT6 (D3H4) | Cell Signaling Technology | RRID:AB_11220421 | #5397 |
| --- | --- | --- | --- |
| ATF3 | Novus | RRID:AB_11014863 | #NBP1-85816 |
| Phospho-STAT6 (Tyr641) | Cell Signaling Technology | RRID:AB_2799514 | #56554 |
| lamin A/C | Cell Signaling Technology | RRID:AB_10545756 | #4777 |
| actin | Sigma-Aldrich | RRID:AB_476693 | #A2066 |
| WASP | Santa Cruz | RRID:AB_628445 | #sc-13139 |
| STAT1 | Cell Signaling Technology | RRID:AB_2198300 | #9172 |
| Phospho-STAT1 (Tyr701) (58D6) | Cell Signaling Technology | RRID:AB_561284 | #9167 |
| Phospho-Cofilin (Ser3) (77G2) | Cell Signaling Technology | RRID:AB_2080597 | #3313 |
| Cofilin (D3F9) | Cell Signaling Technology | RRID:AB_10622000 | #5175 |
| HMGCR | Atlas Antibodies | RRID:AB_2665607 | #AMAb90619 |
| HMGCS1 | Novus | RRID:AB_3295882 | #NBP2-36554 |
| ATP5α (15H4C4) | Abcam | RRID:AB_301447 | #ab14748 |
| HRP AP Donkey Anti-Mouse IgG (H+L) | Jackson Immunoresearch | RRID:AB_2340770 | #715-035-150 |
| HRP AP Donkey Anti-Rabbit IgG (H+L) | Jackson Immunoresearch | RRID:AB_10015282 | #711-035-152 |

**Chemicals & Reagents**

| Chemical/Reagent | Source | Catalog Number |
| --- | --- | --- |
| Cas9 mRNA | Tebu-bio | #L-6125-20 |
| TRI Reagent | Sigma-Aldrich | #T9424 |
| ReliaPrep RNA Cell Miniprep System | Promega | #z6011 |
| SuperScript III Reverse Transcriptase | Thermo Scientific | #18080-044 |
| GoTaq probe qPCR Master Mix | Promega | #A6102 |
| GoTaq qPCR Master Mix | Promega | #A6002 |
| Emerald Master Mix | Takara | #RR330B |
| ROTI®Quant Bradford solution | Carl Roth | #K015.1 |
| Pierce 660nm Protein Assay Reagent | Thrmo Scientific | #22660 |
| Protease inhibitor cocktail tablets, cOmplete | Sigma-Aldrich | #11873580001 |
| Phosphatase inhibitor cocktail tablets, PhosSTOP | Sigma-Aldrich | #04906837001 |
| Mowiol 4-88 | Sigma-Aldrich | #81381-250G |
| DNAse I | Promega | #Z358A |
| ECL Western blotting substrate | Promega | #W1015 |
| DAKO | Agilent | #S3022 |
| Tissue culture reagents and consumables | | |
| rM-CSF | ImmunoTools | #12343113 |
| IL-4 | Peprotech | #214-14 |
| Recombinant Murine RANKL | Peprotech | #315-11 |
| FSL-1 | Invivogen | #tlrl-fsl |
| Pam3CSK4 | Invivogen | #tlrl-pms |
| 25-Hydroxycholesterol (25-HC) | Sigma-Aldrich | #H1015 |
| Lonafarnib | Sigma-Aldrich | #SML1457 |
| GGTI-2133 | Sigma-Aldrich | #G5294 |
| Mycobacterium tuberculosis str. Erdman = ATCC 35801 | Rybniker Lab |  |
| DMEM (1x) + GlutaMAX | Thermo Scientific | #31966-021 |
| DPBS (1x) | Thermo Scientific | #14190-094 |
| Penicillin-Streptomycin (10,000 U/mL) | Thermo Scientific | #15140-122 |
| 0,05% Trypsin-EDTA (1x) | Thermo Scientific | #25300-054 |
| FBS Superior | Biochrom | #S0615 |
| Sodium Pyruvate 100 mM (100x) | Thermo Scientific | #11360-039 |
| TrypLE™ Express Enzym (1x), Phenolrot | Thermo Scientific | #12605036 |
| 15 cm sterile Petri dishes | Greiner | #639161 |
| 6 well plates sterile | Greiner | #657160 |
| Nunc Lab-Tek 8-well chamber glass slides | Thermo Scientific | #177402 |
| Nunc Lab-Tek 8-well chamber Permanox slides | Thermo Scientific | #177445 |
| 35 mm imaging dishes, high glass bottom | Ibidi | #81158 |
| µ-Slide 8 Well Glass Bottom | Ibidi | #80827 |
| CD11b MicroBeads | Miltenyi Biotec | #130-049-601 |
| Chloroform | MERCK | #1.02445.1000 |
| Bovine Serum Albumin (BSA) | Sigma-Aldrich | #A3059 |
| Dimethyl Sulfoxide (DMSO) | Sigma-Aldrich | #20-139 |
| Tween-20 | Sigma-Aldrich | #P1379-500 |
| Triton X-100 | Sigma-Aldrich | #101371900 |
| 2-Mercaptoethanol (14.3M) | Sigma-Aldrich | #M3148-100M |
| SDS | Roth | #CN30.3 |
| HEPES (1 M) | Thermo Scientific | #15630106 |
| PDMS microchannels | 4D Cell | #MC011, #MC019 |

**Buffers**

| Buffer | Components and Final concentrations |
| --- | --- |
| Genotyping Tissue Lysis Buffer | 100 mM Tris/HCl pH 8.5, 5 mM EDTA, 0.2% SDS and 200 mM NaCl |
| TE-buffer | 10 mM Tris-HCl pH 8, 1 mM EDTA pH 8.0 |
| PBS | 137 mM NaCl, 2.7 mM KCl, 10 mM NA_2_PO_4_, 2 mM KH_2_PO_4_ |
| PBST | 1x PBS, 0,5% Triton |
| Blocking buffer for immunostaining | 1x PBS, 3% BSA |
| TBS | 50 mM Tris, 150 mM NaCl, pH 7.5 |
| TBST | 1x TBS, 0.1% Tween-20 |
| Blocking buffer for immunoblotting | 1x TBST, 3% BSA |
| 6x Laemmli buffer | 125 mM Tris-HCl pH 6.8, 20% glycerol, 4% SDS, 10% β-Mercaptoethanol and bromophenol blue |
| RIPA | 50 mM HCl pH 8, 150 mM NaCl, 1% NP-40, 0.5% sodium deoxycholate, 0.1% SDS |
| cytoskeleton-stabilizing lysis buffer | 50 mM PIPES pH 6.9, 50 mM NaCl, 5 mM MgCl_2_, 5 mM EGTA, 0.2 mM dithiothreitol, 0.1% NP-40, 0.1% Tween-20, 5% glycerol, 1 mM ATP and protease inhibitors |
| actin-depolymerizing buffer | 50 mM PIPES pH 6.9, 5 mM MgCl_2_, 10 mM CaCl_2_, 5 µM cytochalasin D |

**Software and Algorithms**

| Software or Algorithms | Source |
| --- | --- |
| Fluoview 2.1c Software | Olympus |
| Photoshop 2025 | Adobe Systems, Inc. |
| Prism 10 | GraphPad Software |
| FIJI | ImageJ (https://fiji.sc/) |
| Huygens Essential 17.10.0p8 64b | Scientific volume imaging |
| ApE plasmid editor | http://jorgensen.biology.utah.edu/wayned/ape/ |
| gRNA design tool | crispr.mit.edu |
| LAS X 3.5.5.19976 | Leica |
| CellPose | http://www.cellpose.org/ |
| ShinyGO 0.82 | http://bioinformatics.sdstate.edu/go/ |
| Mausoleum | http://www.maus-o-leum.de/home.html |
| R-Studio |  |
| Kallisto | https://pachterlab.github.io/ |
| Benchling | https://benchling.com/editor |
| GeneOntology | http://geneontology.org/ |
| Biorender | https://app.biorender.com |
| NIH BIOART | https://bioart.niaid.nih.gov/ |

**Equipment**

| Equipment | Source |
| --- | --- |
| CFX96 Real-Time PCR System | Bio-Rad |
| FV1000 Confocal Microscope | Olympus |
| SZX16 fluorescent stereomicroscopes | Olympus |
| DP72 CCD camera | Olympus |
| ImageQuant LAS4000 reader | GE Healthcare |
| EnSpire Multimode Plate Reader | Perkin Elmer |
| Gel Doc XR+ Imaging System | Bio-Rad |
| NanoDROP 8000 | Thermo Scientific |
| Optima MAX Ultracentrifuge (TLA-55 rotor) | Beckman Coulter |
| UltraView VoX Spinning Disk confocal microscope | Perkin Elmer |
| LSM 980 with Airyscan 2 | Zeiss |
| TCS SP8 confocal microscope | Leica |
| EM 109 transmission electron microscope | Zeiss |
| EVOS FL Auto 2 | Thermo Scientific |
| CASY cell counter & analyzer | OMNI Life Science |
