## Supplementary FIgure S1 for "Atf3 Integrates Lipid and Cytoskeletal Remodeling to Drive Macrophage Fusion"

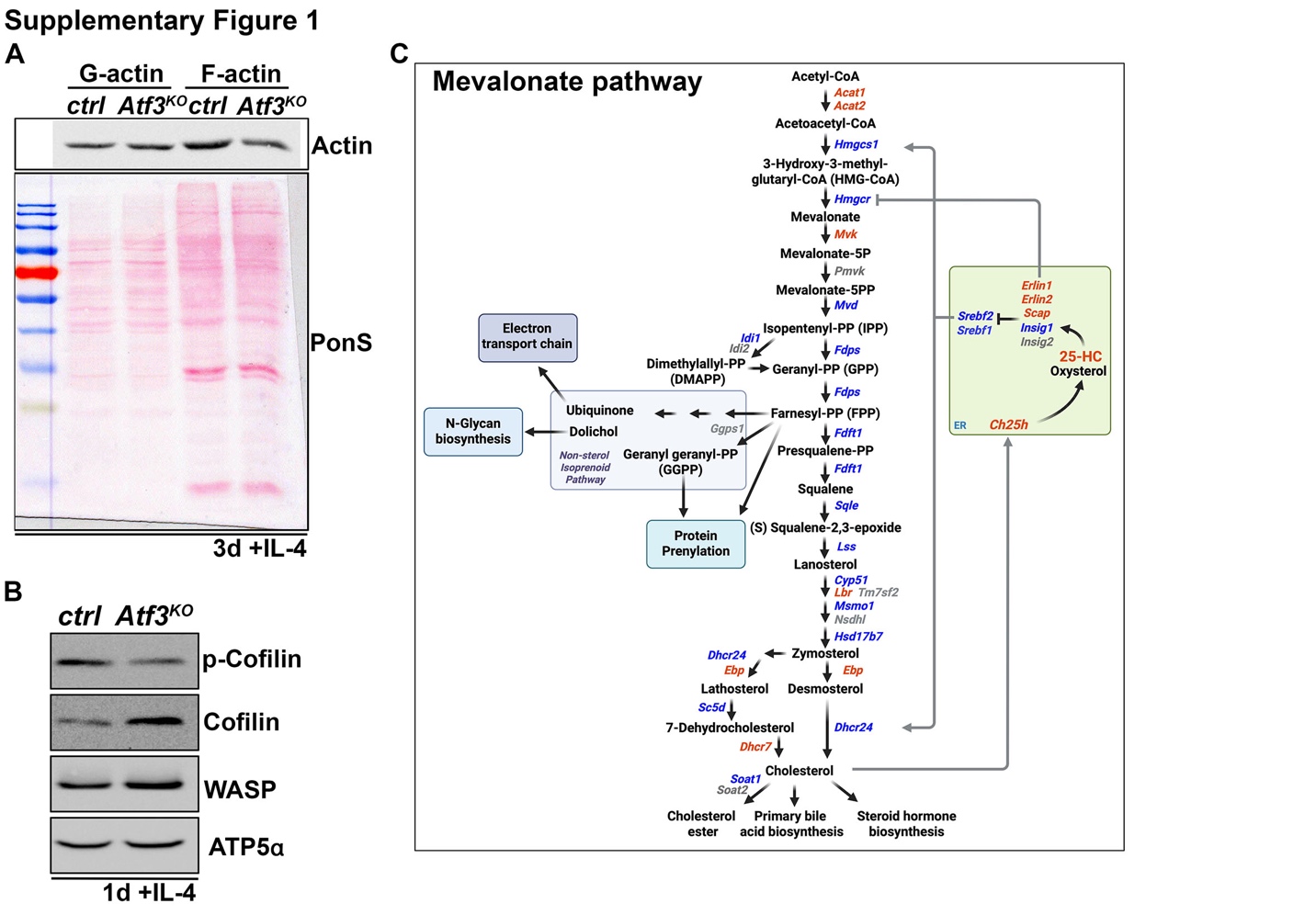

**Supplementary Figure S1. Atf3 loss interferes with actin cytoskeleton remodeling and inhibits the mevalonate pathway.**

**(A-B)** Representative western blots showing decrease of F-actin in *Atf3^KO^* BMDMs compared to *control* following 3-day treatment with IL-4 (A). WASP and total Cofilin protein levels increase in Atf3-deficient IL-4-stimulated BMDMs while phosphorylated inactive form of Cofilin (p-Cofilin) is decreased relative to control (B). Ponceau S (PonS) staining (A) and ATP5α (B) were used as loading control. **(C)** Schematic representation of the mevalonate pathway and ER-resident regulators showing upregulated (red), downregulated (blue), or unchanged (grey) genes in *Atf3^KO^* versus *control* BMDMs after 5 h IL-4 stimulation. Depicted gene regulation is based on normalized RNA-seq counts (n = 4 biological replicates per genotype). Farnesyl pyrophosphate (FPP) is a precursor for cholesterol and non-steroidal isoprenoids, including dolichol, ubiquinone, and substrates for protein prenylation. Cholesterol is further converted to esters, bile acids, steroid hormones, and oxysterols. ER-resident Scap-Insig-Erlin complex senses cholesterol/oxysterol availability controlling processing of SREBP-1 (*Srebf1*) and SREBP-2 (*Srebf2*) transcription factors that regulate expression of genes involved in fatty acid and cholesterol synthesis, respectively. While Erlin and Scap bind cholesterol, Insig binds 25-hydroxycholesterol (25-HC) generated by cholesterol-25-hydroxylase (Ch25h). High cholesterol or 25-HC antagonizes SREBPs processing causing their retention in the ER out of the reach of Golgi located activator proteases. Moreover, 25-HC inhibits the pathway by accelerating proteasome-mediated degradation of Hmgcr. The model was generated using BioRender.com.
