## Supplementary FIgure S2 for "Atf3 Integrates Lipid and Cytoskeletal Remodeling to Drive Macrophage Fusion"

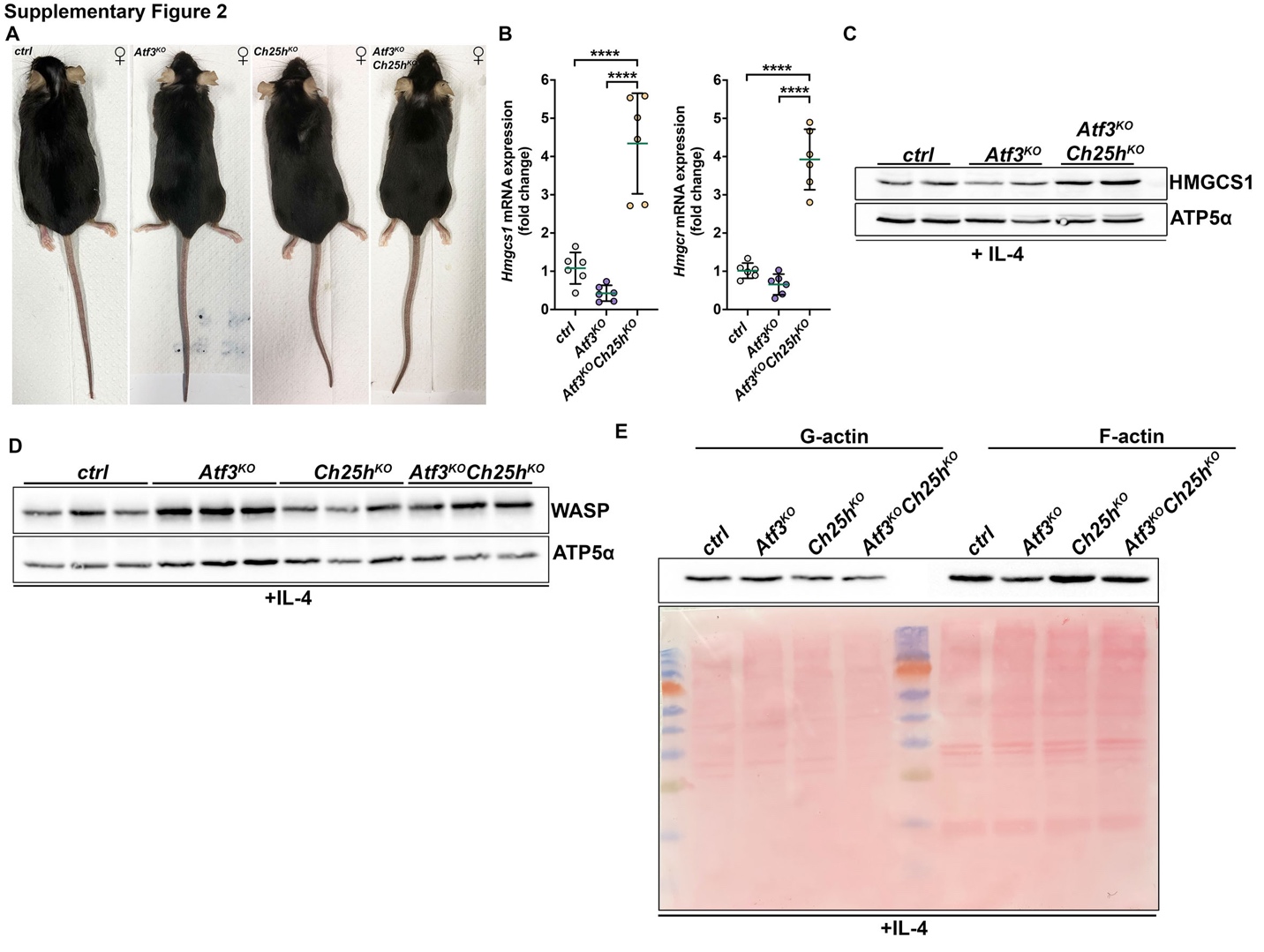
**Supplementary Figure S2. Ch25h loss upregulates mevalonate pathway genes and restores actin polymerization in Atf3-deficient BMDMs.**

**(A)** Representative images of 86-89-weeks old female mice of the indicated genotypes. **(B)** RT-qPCR shows significant upregulation of *Hmgcs1* and *Hmgcr* mRNA expression in *Atf3^KO^Ch25h^KO^* BMDMs compared to *control* and *Atf3^KO^* cells. Data represent means ± s.d.; ***p<0.001, ****p<0.0001, n.s. = non-significant. Statistical significance was determined using ordinary one-way ANOVA with Tukey’s multiple comparison test. Each dot represents an average value from individual animal, n = 6. **(C-D)** Representative western blots showing Hmgcs1 (C) and WASP (D) protein levels in BMDMs of the indicated genotypes. ATP5α was used as loading control. Each genotype is represented by two (C) or three (D) BMDM samples prepared from independent animals. **(E)** Representative western blot showing G- and F-actin levels in IL-4 stimulated BMDMs of the indicated genotypes. The Ponceau S-stained membrane shows equal loading for each of the actin fractions.
